# Structural coordination between active sites of a Cas6-reverse transcriptase-Cas1—Cas2 CRISPR integrase complex

**DOI:** 10.1101/2020.10.18.344481

**Authors:** Joy Y. Wang, Christopher M. Hoel, Basem Al-Shayeb, Jillian F. Banfield, Stephen G. Brohawn, Jennifer A. Doudna

## Abstract

CRISPR-Cas systems provide adaptive immunity in bacteria and archaea by targeting foreign DNA for destruction using CRISPR RNA-guided enzymes. CRISPR immunity begins with integration of foreign sequences into the host CRISPR genomic locus, followed by transcription and maturation of CRISPR RNAs. In a few CRISPR systems, the Cas1 integrase and a Cas6 nuclease are fused to a reverse transcriptase that enables viral sequence acquisition from both DNA and RNA sources. To determine how these components work together, we determined a 3.7 Å resolution cryo-EM structure of a Cas6-RT-Cas1 protein complexed with Cas2, a subunit of the CRISPR integrase. The structure and accompanying mutagenesis experiments provide evidence of bidirectional crosstalk between the Cas1 and RT active sites and unidirectional crosstalk from Cas6 to the Cas1 and RT active sites. Together, these findings suggest regulated structural rearrangements that may coordinate the complex’s different enzymatic activities.

---

CRISPR-Cas (clustered regularly interspaced short palindromic repeats-CRISPR associated) systems provide adaptive immunity against viruses for bacteria and archaea^1,2^. Immunity begins when an integrase comprising the conserved Cas1 and Cas2 proteins incorporates segments of foreign DNA – protospacers – into the host CRISPR locus, which consists of direct repeats separated by virally-derived spacers^3–5^. Transcription and transcript processing generate CRISPR RNAs (crRNAs) that assemble with effector complexes to identify and destroy foreign nucleic acids bearing sequence complementarity to the ~20-nucleotide (nt) spacer segment of the crRNA^6–11^.

Although most CRISPR spacer sequences are derived from DNA phage, a minority of CRISPR-Cas systems include a reverse transcriptase (RT), enabling spacer acquisition from RNA sources^20–29^. In some cases, the RT occurs as a fusion protein together with Cas1 on the RT’s C-terminus that can facilitate DNA-based storage of transcriptional information and has been used as a tool to record cellular transcriptional activity^29^. RT-Cas1s occur primarily in type III CRISPR-Cas systems that target both RNA and DNA, raising the interesting possibility that these variants provide adaptive immunity against RNA phage or other RNA elements^8,30–33^. Notably, many of these RT-Cas1 fusion proteins include an N-terminal Cas6 domain^24,26,34^, which processes CRISPR transcripts into mature crRNAs^35–37^. A recent study showed that the Cas6 domain of a Cas6-RT-Cas1 fusion is required for RNA spacer acquisition and co-evolves with the RT domain^34^. However, the molecular basis for functional coordination between Cas1, RT and Cas6 remains unknown.

To determine how RT activity might coordinate with the CRISPR integrase and with the Cas6 maturation nuclease in the context of a Cas6-RT-Cas1 fusion protein, we determined the 3.7 Å resolution structure of a naturally-occurring *Thiomicrospira* Cas6-RT-Cas1—Cas2 complex using single particle cryo-electron microscopy (cryo-EM) and investigated its activities *in vitro*. We show that in spite of a significant difference in arrangement of the Cas1—Cas2 disposition compared to the most extensively studied CRISPR integrase, the *E. coli* integrase, the Cas6-RT-Cas1—Cas2 complex is functional in carrying out site-specific full-site CRISPR spacer integration of double-stranded DNA (dsDNA) substrates and providing length selectivity. The complex is capable of catalyzing complementary DNA (cDNA) synthesis with both RNA and DNA templates with minimal homology requirements between primer and template, though the structure reveals a unique RT conformation compared to other RT structures. Finally, the structural data and experimental mutagenesis indicate strong crosstalk between the three domains - most notably, an RT-helix that plays a significant role in both RT and Cas1 activities and may be involved in a large structural rearrangement, providing further insight into how the complex coordinates its different activities.

## RESULTS

### Architecture of the Cas6-RT-Cas1—Cas2 complex

To investigate the structure of a Cas6-RT-Cas1—Cas2 complex and its activities, we purified a Cas6-RT-Cas1 and its associated Cas2 protein, which are part of a newly identified *Thiomicrospira* type III CRISPR locus (Fig. 1a). To form the complex, the Cas6-RT-Cas1 and Cas2 proteins were assembled with a DNA substrate designed to mimic a genomic integration intermediate^16^ (Fig. 1a,b, Supplementary Fig. 1a). The structure of the resulting Cas6-RT-Cas1—Cas2 complex was determined by single particle cryo-EM at an overall resolution of 3.7 Å (Fig. 1b-d, Supplementary Fig. 1b, 2a-h). Approximately half of the complex displayed relatively lower local resolution, and refinement using a mask that excluded this region generated a reconstruction at 3.4 Å resolution with improved interpretability of high-resolution features. A model corresponding to two Cas2 chains and portions of four Cas6-RT-Cas1 chains was *de novo* built and refined into this masked reconstruction of the partial complex. Portions of three Cas6-RT-Cas1 chains that were excluded by the mask were then rigid body docked and refined in the full complex reconstruction. The final model consists of two entire Cas2 chains, two entire Cas6-RT-Cas1 chains (excluding 5 poorly resolved loops), and the Cas1 domain from two additional Cas6-RT-Cas1 chains.

**Figure 1.**
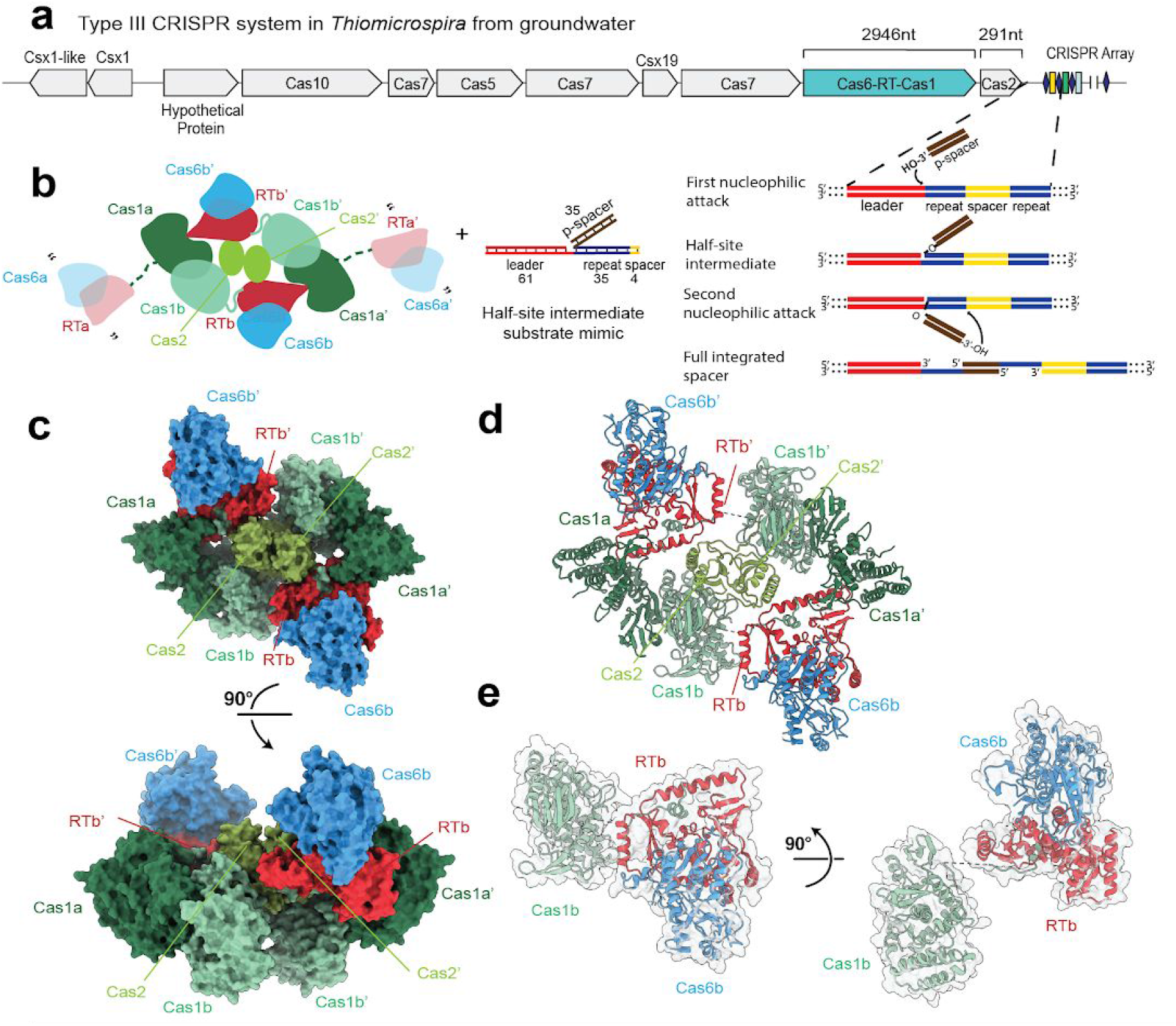
A natural Cas6-RT-Cas1 fusion protein within a type III CRISPR-Cas system. **a)** Illustration of the CRISPR genomic locus encoding the proteins used in this study and schematic of the CRISPR spacer integration reaction. Red, leader; blue, repeat; yellow, spacer; brown, dsDNA protospacer. **b)** Cartoon showing Cas6-RT-Cas1 and Cas2 proteins and the DNA substrate, mimicking the half-site intermediate depicted in **a**, used in complex assembly. Proteins colored by domain: Cas6, blue; RT, red, Cas1a, dark green; Cas1b, turquoise; Cas2, lime green. Visible linkers RT-Cas1 linkers are shown as solid turquoise lines. Missing Cas6-RT domains (Cas6a/RTa and Cas6a’/RTa’) that are present in the complex but not visible are depicted as semi-transparent, connected to the rest of the structure by dashed dark green lines. Quotation marks indicate presumed mobility. Lengths of the DNA substrate strands are indicated. **c)** Overall structure of the complex depicted in surface representation and 90° rotation, with color-coding as shown in **b**. **d)** Overall structure of the complex, depicted in ribbon representation. **e)** Architecture of a single Cas6-RT-Cas1 protomer and 90° rotation.

As observed for previous CRISPR integrase structures^16,18,19^, the Cas6-RT-Cas1—Cas2 complex is heterohexameric, consisting of a central Cas2 dimer and two distal Cas6-RT-Cas1 dimers (Fig. 1b-d). The heterohexameric architecture of the core Cas1—Cas2 is a hallmark of CRISPR integrases and is critical to their internal ruler mechanism dictating the length of the spacers that are integrated and their ability to catalyze full-site integration^16,18,19^. Extending from the Cas1—Cas2 core are the Cas6-RT lobes, which are positioned orthogonal to the Cas1 domains relative to the central Cas2 dimer. The Cas6-RT lobe is connected to its associated Cas1 domain via a flexible linker, with the RT domain proximal to the central Cas2 dimer and the Cas6 domain closely abutting the RT domain (Fig. 1c-e). We observed density for the Cas6-RT lobe in two of the four Cas6-RT-Cas1 protomers (Cas1b and b’), presumably due to flexibility of the Cas6-RT domains in the other two Cas6-RT-Cas1 protomers (Cas1a and a’). No density was observed for the DNA substrate despite co-migration during size-exclusion chromatography (Supplementary Fig. 1a).

The Cas1 and Cas2 domains largely resemble those of the *E. coli* Cas1—Cas2 complex; however, there are some notable differences. When compared to the *E. coli* Cas1—Cas2 complex structure, the C-terminal α-helical domains of a given Cas1 are very similar (2.3 Å RMSD), while the linker and N-terminal β-sheet domains are offset by ~5-7 Å (Supplementary Fig. 3a,b). Each Cas2 protomer has a ferredoxin-like fold, as observed in previous Cas2 structures^38–40^. However, compared to the *E. coli* Cas1—Cas2 complex^12^, the β-sheet in Cas2 is formed by fewer β-strands and the C-terminal domain crosses the central axis to pack against the face of the opposing subunit, resulting in an altered dimer interface. Additionally, the C-terminal segment (comprising amino acids 88-96) interacts with Cas1b rather than with Cas1b’ as in the *E. coli* Cas1—Cas2 structure.

Previous studies have shown that CRISPR-associated RTs are most closely related to the RTs encoded by group II introns^24–27^. We observe that the RT domain shares the characteristic palm and fingers regions of the canonical right-handed fold in other RT structures but is missing the thumb/X subdomain that is present in retroviral and group II intron RTs^41–44^. Compared to the group II intron RT structure^41^, the finger domains are very similar with an overall RMSD of 1.9 Å, while the palm domains are strikingly different. Interestingly, in the present structure, the palm domain is reoriented such that the FADD motif, containing active site residues conserved across diverse RT subtypes, points away from the substrate binding cleft, suggesting that this conformation may not be competent for reverse transcription.

Finally, the structure of the Cas6 domain closely resembles those of previous Cas6 structures^34,36,45–48^ (Supplementary Fig. 3c). When compared directly to the structure of the Cas6 domain from the *Marinomonas mediterranea* (MMB-1) Cas6-RT-Cas1 protein^34^, the two structures align with RMSD of 2.4 Å, with the greatest variance in the β-strand of the C-terminal RNA recognition motif (RRM) fold, perhaps due to the subsequent connection to the RT domain, absent in the MMB-1 Cas6 structure (Supplementary Fig. 3d).

### Substrate preferences of Cas6-RT-Cas1—Cas2 for cleavage-ligation reactions

Previous biochemical and genomic experiments demonstrated both RT and integration activities for the MMB-1 RT-Cas1 fusion protein. To test the integrase function of the *Thiomicrospira* Cas6-RT-Cas1—Cas2 complex, we first investigated its substrate preferences for integration into a target DNA molecule and the structural elements that may be involved in the cleavage-ligation reactions. We conducted integration assays by incubating the purified Cas6-RT-Cas1 and Cas2 proteins with a 5’-fluorophore-labeled 35-base pair (bp) dsDNA or a 35-nt single-stranded DNA (ssDNA) or single-stranded RNA (ssRNA) substrate and a supercoiled target plasmid containing the CRISPR locus. Deoxynucleoside triphosphates (dNTPs) were supplied to reactions with RNA substrates. We observed protospacer ligation by examining the incorporation of the fluorophore into the target DNA molecule (Fig. 2a). The results show that the Cas6-RT-Cas1—Cas2 complex can catalyze ligation of all three substrates into the target plasmid (Fig. 2b). Interestingly, the Cas6-RT-Cas1 protein alone is able to catalyze protospacer ligation with DNA substrates. While Cas2 alone shows no activity, its presence increases the cleavage-ligation efficiency of the complex. Time-course experiments that quantified the extent of plasmid-nicking in each reaction showed 100% and 78% conversion from supercoiled plasmid target to open-circle product with dsDNA and ssDNA, respectively, 25% with ssRNA, and 13% with no protospacer over two hours (Fig. 2c, Supplementary Fig. 4a). These results show that this *Thiomicrospira* Cas6-RT-Cas1—Cas2 integrase is significantly more efficient with DNA substrates than RNA substrates for cleavage-ligation.

**Figure 2.**
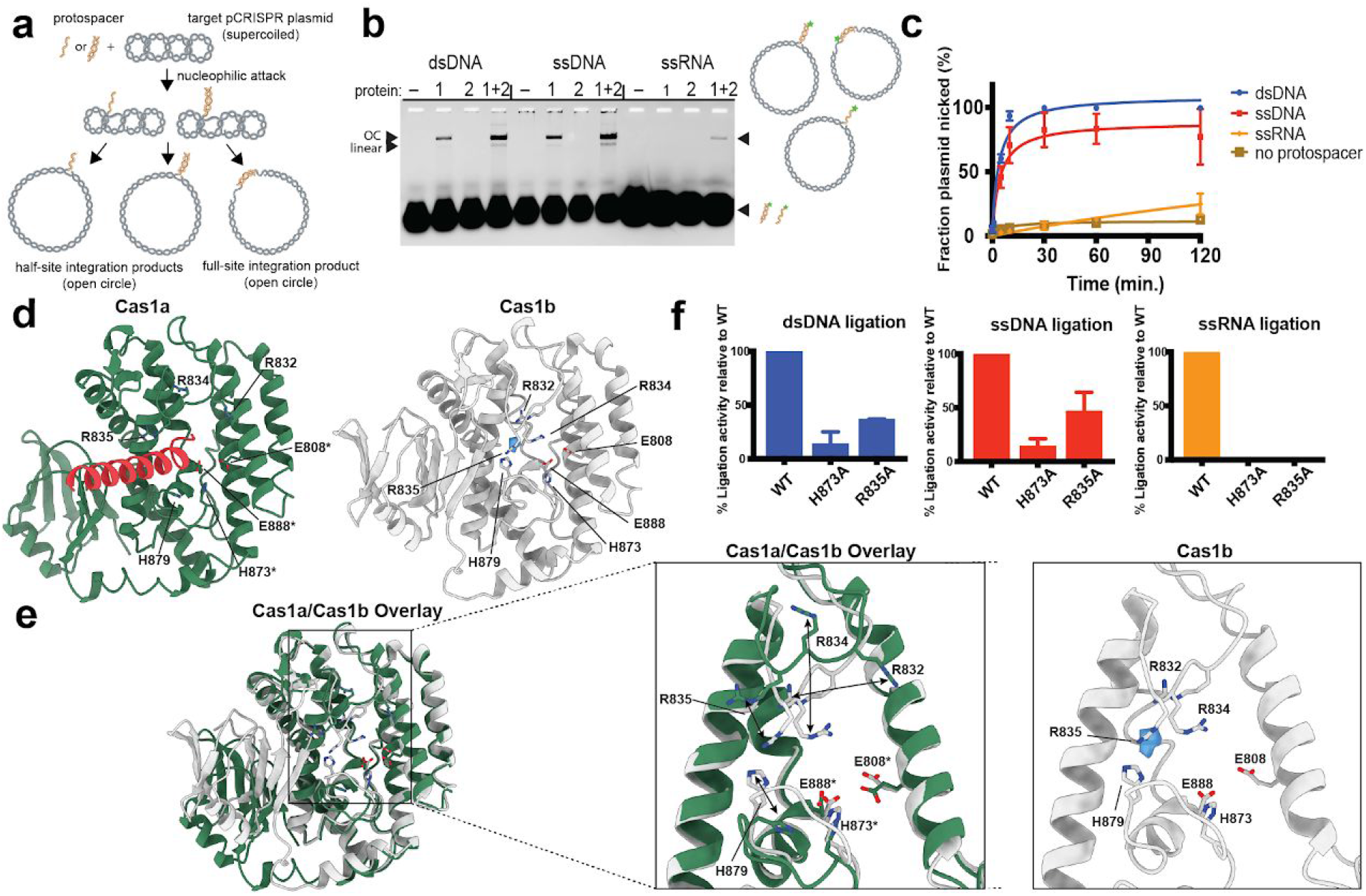
Cas6-RT-Cas1—Cas2 catalyzes cleavage-ligation reactions with dsDNA, ssDNA, and ssRNA substrates *in vitro*. **a)** Schematic of *in vitro* integration reaction of DNA or RNA protospacer into target supercoiled pCRISPR plasmid. **b)** Integration assay with fluorescent 35-nt dsDNA (0.5 μM), ssDNA (0.5 μM), and ssRNA (4 μM) protospacers. Open-circle (OC) and linear ligation products are indicated. Star indicates 6-carboxyfluorescein label. **c)** Time-course integration assay comparing extent of plasmid-nicking with dsDNA, ssDNA, and ssRNA protospacers (35-nt; 4 μM). Fraction plasmid nicked is calculated as the fraction of open-circle products relative to all plasmid (mean ± sd, *n* = 3 independent experiments). Experimental fits are shown as solid lines. Representative gels are shown in Supplementary Fig. 4a. **d)** Comparison of Cas1a (dark green) and Cas1b (white), with RT-helix (red) and unknown density (blue). Active site and surrounding residues are shown in stick configuration. Active site residues are labeled with an asterisk *. **e)** Cas1a and Cas1b overlay and closeup of active site. Arrows indicate large conformational differences. **f)** Ligation of 35-nt fluorescent dsDNA (0.5 μM), ssDNA (0.5 μM), and ssRNA (4 μM) protospacers into target pCRISPR by mutant Cas6-RT-Cas1—Cas2 proteins. Percent ligation activity is calculated as the fraction of fluorescent products from the mutant Cas6-RT-Cas1—Cas2 relative to that from the WT (mean ± sd, *n* = 3 independent experiments). Representative gels are shown in Supplementary Fig. 7c. Source data for panels **c** and **f** are available in Supplementary Data Set 3. Uncropped gels are available in Supplementary Data Set 2.

We wondered if there are unique elements near the Cas1 active site that enable RT-associated CRISPR integrases to catalyze cleavage-ligation chemistry with RNA substrates, a process not observed for non-RT-associated Cas1s. We found a cluster of residues (R832, R834, R835, and H879) adjacent to the active site that undergoes a large conformational change between Cas1a and Cas1b. In the inactive Cas1b domains, these residues appear to coordinate an unknown density, potentially corresponding to a metal ion or water molecule (Fig. 2d,e). In the catalytic Cas1a domains, this density is not observed and the residues of interest are pushed away from each other, seemingly to accommodate the insertion of an α-helix from the RT domain. Sequence alignments across a diverse sample of RT-associated Cas1 variants show that there is ~100%, ~60%, ~70%, and ~35% conservation for R832, R834, R835, and H879, respectively, across RT-associated Cas1s (Supplementary Fig. 4b). The latter three residues show a slightly lower sequence conservation across non-RT-associated Cas1s and are not conserved in the *E. coli* Cas1 (Supplementary Fig. 3a,b). We hypothesize that this cluster of residues may play an important role for the Cas6-RT-Cas1’s functions.

To examine the potential role of these residues in protospacer ligation, we conducted integration assays with the R835A mutant and the Cas1 active site mutant H873A. We quantified the relative amounts of protospacer ligation to parallel wild-type (WT) controls. As expected, the H873A mutant abolishes almost all protospacer ligation (Fig. 2f). Interestingly, the R835A mutant also has significant effects, with a 65% and 55% reduction for dsDNA and ssDNA ligation and >99% reduction for ssRNA. Together, these findings identify R835 as a potential structural element that contributes toward Cas6-RT-Cas1—Cas2’s ability to accommodate RNA substrates for protospacer ligation.

To determine whether the *Thiomicrospira* Cas6-RT-Cas1—Cas2 complex can catalyze cleavage-ligation in a site-specific manner with all three substrates, we conducted integration reactions using a short dsDNA linear target molecule comprising 49 bp of the leader sequence, the 35-bp repeat, and 15 bp of the adjacent spacer (Supplementary Fig. 5a). Although the results show many off-target ligation products, the predominant product corresponds to cleavage-ligation at the spacer end of the repeat for all three protospacers (Supplementary Fig. 5b,c). Curiously, while previous work on the MMB-1 system suggests that the cleavage-ligation products are potential substrates for target-primed reverse transcription^27,49,50^, we do not observe any bands that represent extension of the 3’ end of the DNA after cleavage-ligation when dNTPs are supplied (Supplementary Fig. 5c). Furthermore, unlike what is observed for the MMB-1 system, dNTPs are not required for RNA ligation. The similar pattern of ligation products for all three substrates suggests that the mode for target recognition is likely the same with both RNA and DNA protospacers, although other factors likely contribute to its difference in efficiency.

### Cas6-RT-Cas1—Cas2 exhibits length selectivity for ssDNA and dsDNA substrates

At the heart of the Cas6-RT-Cas1—Cas2 structure lies the heterohexameric Cas1—Cas2 core, which has some interesting differences from the *E. coli* integrase structure, raising questions about its functions as a CRISPR integrase. Two positively charged regions of the *E. coli* Cas1—Cas2 structure are critical for protospacer substrate binding: the “Arginine Channel,” which contacts the DNA substrate where the duplex terminates and the single-stranded overhang enters the active site, and the “Arginine Clamp,” which stabilizes the middle of the duplex^18^. Although similar charged regions occur in the Cas1 and Cas2 dimers of the Cas6-RT-Cas1—Cas2 structure, the Cas2s and Cas1a’/Cas1b’ dimer are rotated further away from the Cas1a/Cas1b such that these charged regions are no longer in the same linear plane as observed in the *E. coli* Cas1—Cas2 structures^16,18,51^ (Fig. 3a,b). The differences in the arrangement of the Cas1—Cas2 components relative to their observed positioning in the *E. coli* Cas1—Cas2 integrase led us to test whether the Cas6-RT-Cas1—Cas2 complex retains the intrinsic ruler mechanism characteristic of CRISPR integrases. Results of integration assays performed with 15- to 115-nt ssDNA and dsDNA substrates showed that the complex has some length selectivity for ssDNA and dsDNA. It is most active with 15- to 55-nt substrates and its activity decreases with substrates longer than 55 nt (Fig. 3c,d). This distribution is larger than the range of spacers observed in the CRISPR array (30-47 nt), though it is not surprising since half-site reactions tend to accept a larger variety of substrate lengths than full-site reactions^4^. What is more surprising is that the length distribution is similar for ssDNA and dsDNA protospacers, suggesting that the complex is able to narrow the selection of protospacer lengths in a manner that does not depend on interaction with a DNA duplex or access to two 3’ nucleophilic ends.

**Figure 3.**
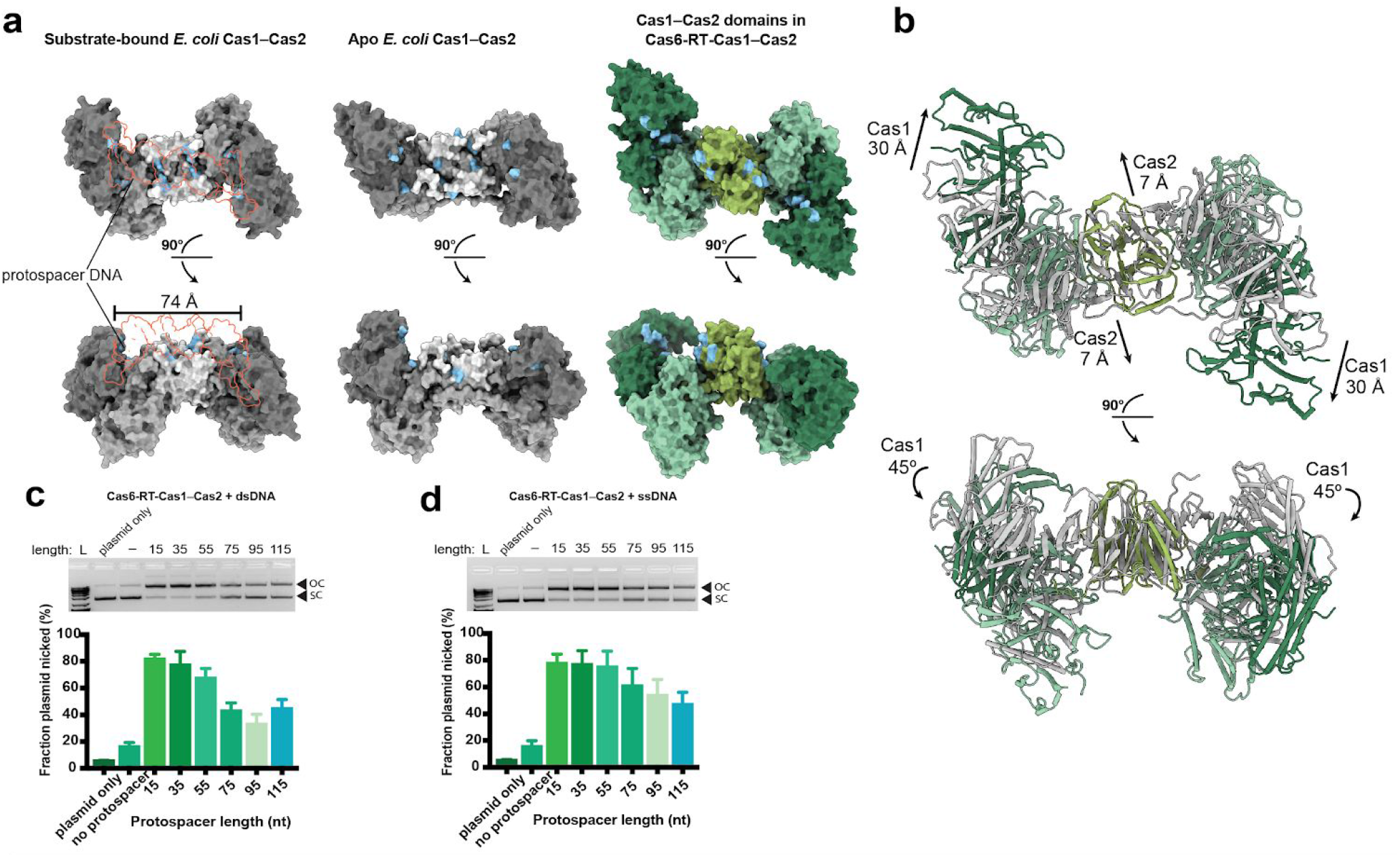
Cas6-RT-Cas1—Cas2 shows length selectivity for dsDNA and ssDNA substrates for integration. **a)** Comparison of Cas1—Cas2 disposition within the Cas6-RT-Cas1—Cas2 complex (color-coding from Fig. 1) to substrate-bound *E. coli* Cas1—Cas2 integrase (PDB:5DS5) and apo *E. coli* Cas1—Cas2 integrase (PDB:4P6I) (Cas1a, dark gray; Cas1b, medium gray; Cas2, light gray), shown in surface representation. Two views are shown related by a 90° rotation. Arginine clamp and arginine channel residues are colored in blue. Protospacer substrate is depicted in red outline. **b)** Overlay of Cas1—Cas2 domains of Cas6-RT-Cas1—Cas2 complex (color-coding from Fig. 1) with apo *E. coli* Cas1—Cas2 integrase (PDB:4P6I) (gray), aligned via Cas2 dimer, shown in ribbon representation. Two views are shown related by a 90° rotation. Arrows indicate conformational difference between Cas1—Cas2 domain of Cas6-RT-Cas1—Cas2 complex and that of *E. coli* Cas1—Cas2. **c)** Cas6-RT-Cas1—Cas2 integration assays with variable length dsDNA protospacers (15 to 115 bp). Supercoiled plasmid target (SC) and open-circle products (OC) are indicated. Fraction plasmid nicked is calculated as the fraction of open-circle products relative to all plasmid (mean ± sd, *n* = 3 independent experiments). **d)** Cas6-RT-Cas1—Cas2 integration with variable length ssDNA protospacers (15 to 115 nt), quantifying fraction plasmid nicked (mean ± sd, *n* = 3 independent experiments). Source data for panels **c** and **d** are available in Supplementary Data Set 3.

### Cas6-RT-Cas1—Cas2 catalyzes full-site integration of dsDNA substrates

We next set out to determine whether the Cas6-RT-Cas1—Cas2 complex can catalyze full-site CRISPR spacer integration. We devised an assay using chloramphenicol resistance as a selection marker for plasmids with protospacer insertion events near the target region^52,53^. The reporter construct was designed with 163 nt of the leader sequence and the CRISPR repeat sequence immediately upstream of a chloramphenicol resistance gene with a missing ribosomal binding site (RBS) sequence and start codon (Fig. 4a). Insertion of a spacer supplying the missing RBS sequence and start codon to the target region allows translation of the chloramphenicol resistance gene transcript. We performed *in vitro* integration assays with this reporter plasmid and a protospacer with an RBS sequence and start codon, transformed the integration products into *E. coli*, plated the transformants on agar containing chloramphenicol, and sequenced the surviving colonies.

**Figure 4.**
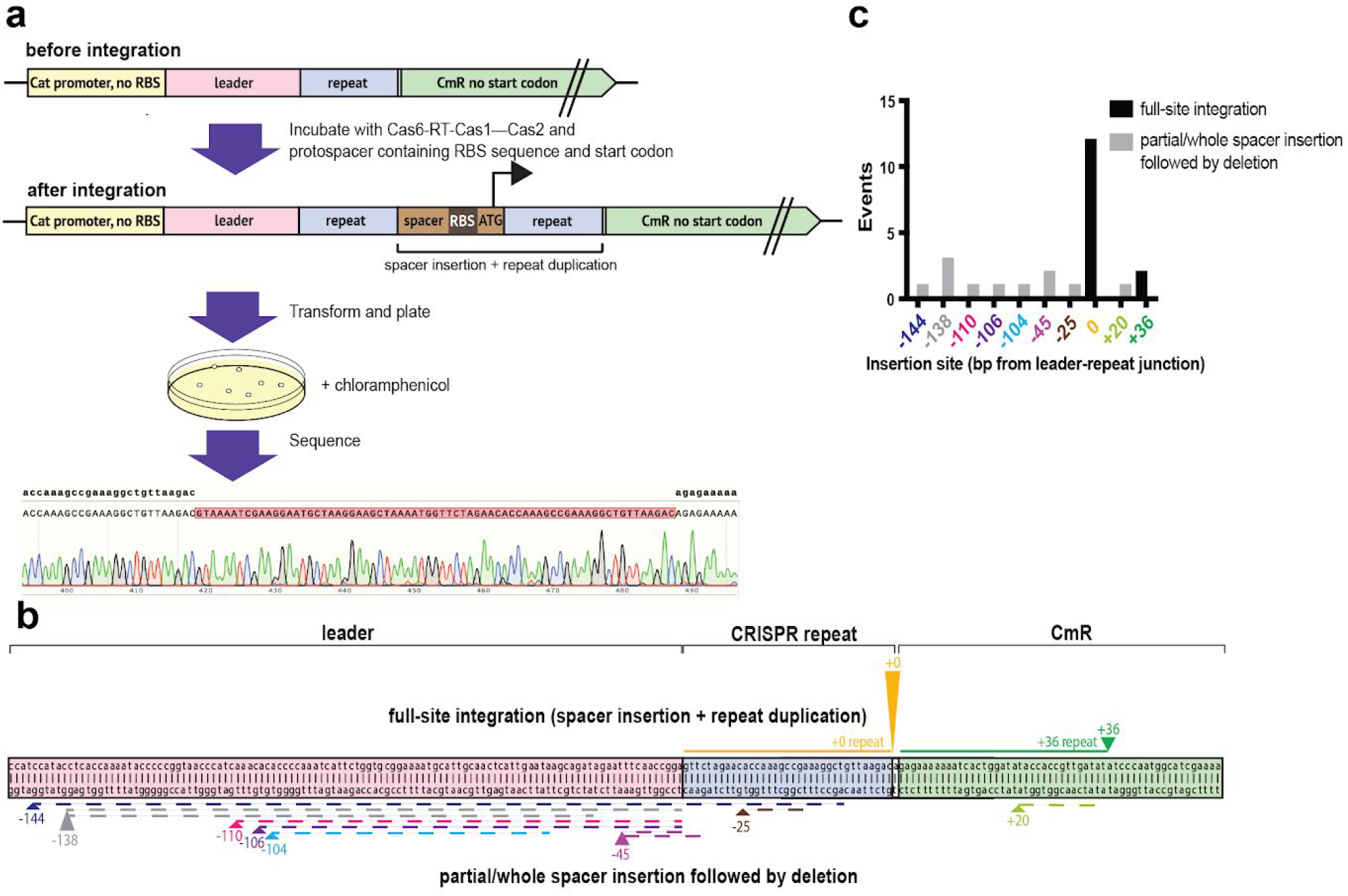
Cas6-RT-Cas1—Cas2 conducts site-specific full-site integration. **a)** Schematic of chloramphenicol selection screen for full-site integration events near the leader-repeat junction. The selection plasmid contains a CRISPR leader and repeat upstream of a chloramphenicol resistance gene (*CmR*) with the RBS and start codon removed. Full-site integration of a protospacer containing an RBS and start codon allows for translation of the *CmR*. Transformants are plated on chloramphenicol plates and clones are sequenced using Sanger sequencing. **b)** Representation depicting spacer insertion events near the leader-repeat junction in a selection plasmid with a 163 bp leader. The arrowheads indicate the insertion sites and the height is scaled to the number of spacer insertion events. The arrows on top indicate full-site integration events, with the corresponding solid colored line adjacent to the arrowhead representing the sequence that is duplicated after spacer insertion. The arrows on the bottom indicate partial/whole spacer insertion events followed by a deletion, with the corresponding dashed colored line adjacent to the arrowhead representing the sequence that is deleted with the spacer insertion. **c)** Number of spacer insertion events near the leader-repeat junction. Insertion sites follow color-coding in **b**. The black bars represent full-site integration events with a 35 bp repeat duplication of the adjacent sequence and the gray bars represent partial/whole spacer insertion followed by a deletion of variable length. Source data for panel **c** are available in Supplementary Data Set 3.

The results show that the Cas6-RT-Cas1—Cas2 complex catalyzes full-site integration with dsDNA substrates. Full-site integration events, characterized by spacer insertion followed by duplication of the adjacent 35 bp, represent the majority of the insertion events (14 out of 25) (Fig 4b,c). The other insertion events appear to be the result of incomplete or abortive integration, characterized by partial/whole spacer insertions followed by a deletion of variable length of the adjacent sequence. While the incomplete/abortive integration events occurred at many off-target locations, full-site integration events were highly specific, with 12 out of the 14 full-site integration sites located at the leader-repeat junction. This suggests that while the complex might attempt integration at off-target sites, it maintains specificity for full-site integration. We did not observe full-site integration with ssDNA or ssRNA protospacers. It is possible that the Cas6-RT-Cas1—Cas2 complex alone can only conduct half-site reactions with single-stranded protospacers *in vitro* and other host factors may be necessary to support full-site integration with those substrates.

### Unique RT conformation results in close contact with Cas1 active site

Although the RT domain has been implicated in RNA-derived CRISPR spacer integration^27^, its mechanism and possible coordination with Cas1 is not known. Similar to other RT structures^41–44^, the RT active site of the Cas6-RT-Cas1—Cas2 complex resides in the palm region and consists of three conserved aspartate residues located on a three-stranded antiparallel β-sheet (Fig. 5a-c). Structural alignment with the group II intron RT reveals that despite close alignment in the fingers region, the palm region of the RT domain is dramatically offset, resulting in a ~90° rotation of the β-sheet containing the active site^41^ (Fig. 5b,c). Possibly as a result, the two active site aspartates between β6 and β7 are further removed from the third active site aspartate between β5 and α5. This rotation is surprising given that there is comparatively little variation among the group II intron RT and other retroviral RT active sites^41–44^, raising the possibility that the current conformation may represent an inactive conformation.

**Figure 5.**
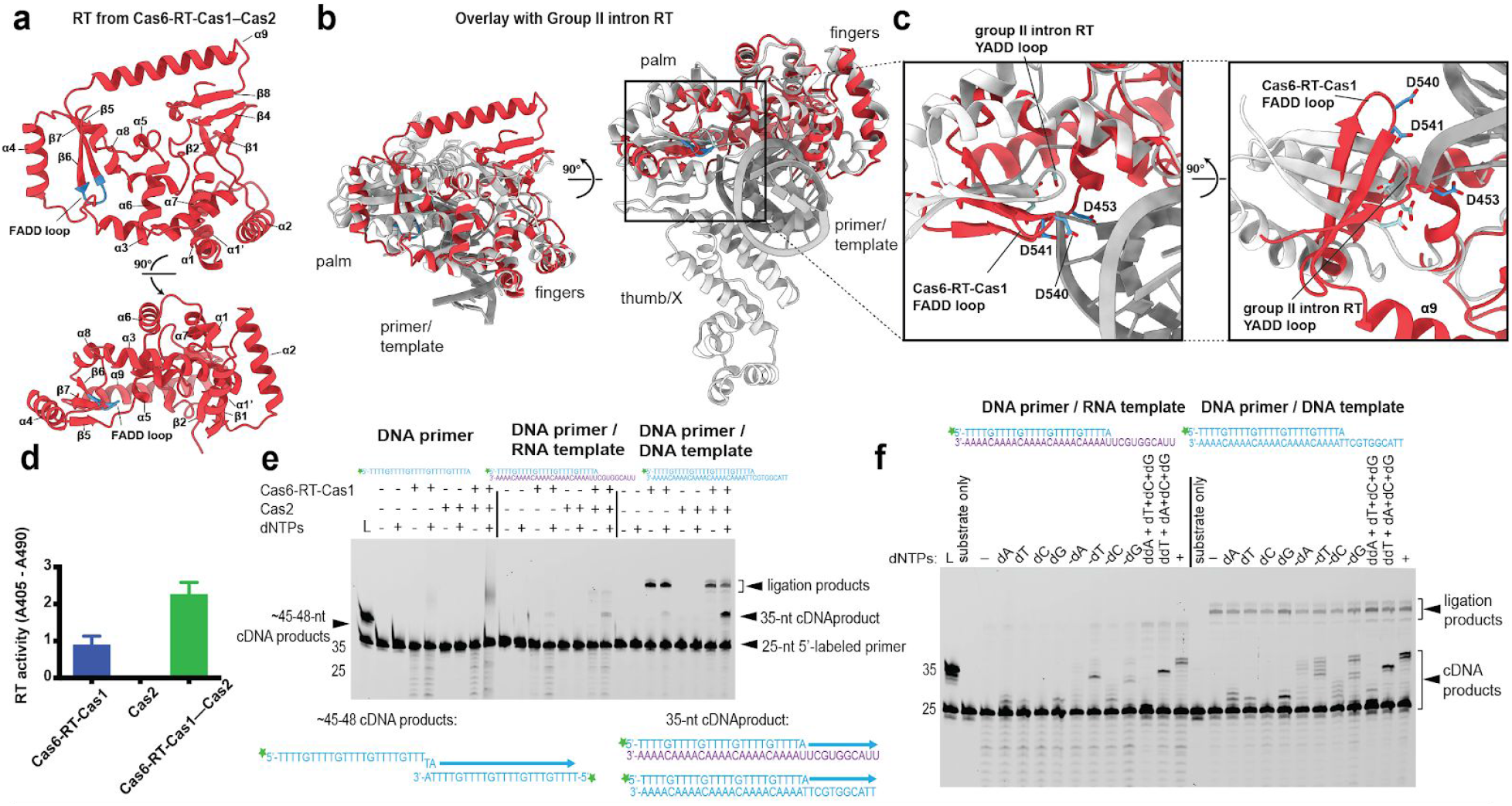
Cas6-RT-Cas1 catalyzes cDNA synthesis with both RNA and DNA templates. **a)** Architecture of RT domain from Cas6-RT-Cas1—Cas2 (red) and 90° rotation, with FADD motif (blue) indicated. **b)** Alignment and overlay of RT domain from Cas6-RT-Cas1—Cas2 (red) with group II intron RT (white) bound to primer/template substrate (PDB: 6AR1) and 90° rotation, FADD motif from Cas6-RT-Cas1—Cas2 colored blue and YADD motif from group II intron RT colored light blue. Palm, fingers, and thumb/X regions and primer/template are indicated. **c)** Closeup of RT active site residues of Cas6-RT-Cas1—Cas2 (blue) and group II intron RT (light blue), shown in stick configuration. Two views are shown related by a 90° rotation. **d)** RT activity assays comparing Cas6-RT-Cas1 alone and Cas6-RT-Cas1—Cas2. RT activity is measured using an ELISA-based colorimetric reverse transcriptase activity assay (mean ± sd, *n* = 3 independent experiments) (Catalog No. 11468120910, Roche Diagnostics, Indianapolis, IN). **e)** Template-driven cDNA synthesis reactions off a fluorescent DNA primer annealed to DNA and RNA templates. Expected cDNA synthesis reactions are indicated. Star indicates 6-carboxyfluorescein label. **f)** Template-driven cDNA synthesis reactions in the absence of different dNTPs and in the presence of added ddNTPs. Uncropped gels are available in Supplementary Data Set 2.

In solution, however, Cas6-RT-Cas1 alone catalyzes reverse transcription, an activity that is enhanced upon addition of Cas2 (Fig. 5d). Using substrates consisting of a 25-nt labeled DNA primer annealed to a 35-nt DNA or RNA template, we observe extension of the primer from 25 to 35 nt, corresponding to the full template length, when dNTPs are supplied (Fig. 5e). There are some unexpected dNTP-independent products around 50-55 nt that do not form when the Cas1 active site mutant is used, suggesting that they are ligation products (Fig. 5e,f, Supplementary Fig. 6a). Interestingly, in the lane that has only the DNA primer as a substrate, a smeary dNTP-dependent band forms around 45-48 nt, which matches the length of a cDNA product that forms when a second copy of the 25-nt DNA substrate is used as a template for primer extension with only 1-2 bp homology. A followup experiment using a 5’-labeled 35-nt DNA or RNA oligonucleotide with the original template sequence also resulted in template-dependent extension of the substrate with only 1 bp homology (Supplementary Fig. 6b). These results suggest that the Cas6-RT-Cas1—Cas2 complex is able to catalyze primer extension with DNA and RNA templates with minimal homology requirements.

To examine RT fidelity, we conducted dNTP drop-out experiments and supplied specific dideoxynucleoside triphosphates (ddNTPs) to terminate primer extension. The results show that Cas6-RT-Cas1—Cas2 complex is faithful to the RNA template, terminating cDNA synthesis at the correct position along the sequence where a necessary dNTP is missing or after a ddNTP is incorporated (Fig. 5f). The complex is slightly more error-prone with the DNA template with bands suggesting some misincorporation of additional dNTPs. The results demonstrate that while the visible RT conformation is likely inactive, the complex can carry out primer extension with both RNA and DNA templates, with slightly higher fidelity for RNA than DNA.

### Structural elements contribute to crosstalk between Cas6, RT, and Cas1 active sites

The structure reveals intriguing elements suggesting potential interplay between the Cas1, RT, and Cas6 domains. The most notable is an RT α-helix that serves as a direct connection between the RT and Cas1 active sites. While the N-terminal end of the RT-helix α9 is attached to the β-sheet containing the RT active site, the C-terminal end points into the catalytic Cas1a active site center (Fig. 6a,b). The helix insertion may be directly related to the conformational difference between Cas1a and Cas1b shown earlier. Structural alignments with other substrate-bound Cas1—Cas2 structures suggest that the helix insertion would result in considerable steric obstruction for the protospacer and target, perhaps even blocking access to the Cas1 active site entirely (Supplementary Fig. 7a,b). The RT-helix would likely need to pull out in order for Cas1 to catalyze integration. We hypothesize that this RT-helix motion may be associated with the rotation of the connected β-sheet, which may bring the RT active site closer to the canonical conformation seen in other RT structures^41–44^. A closer look at the region around the RT-helix reveals additional clues on how this movement could potentially be regulated. The flexible RT-Cas1 linker wraps around the RT-helix, raising the possibility of its involvement in aiding the helix’s movement. The RT-helix also interacts with Cas2, suggesting that rearrangement of the Cas2s could also impact the position of the RT-helix and the attached β-sheet containing the RT active site. An examination of continuous heterogeneity within the cryo-EM data with 3D variability analysis in cryoSPARC is consistent with complex motion in this region (Supplementary Video 1).

**Figure 6.**
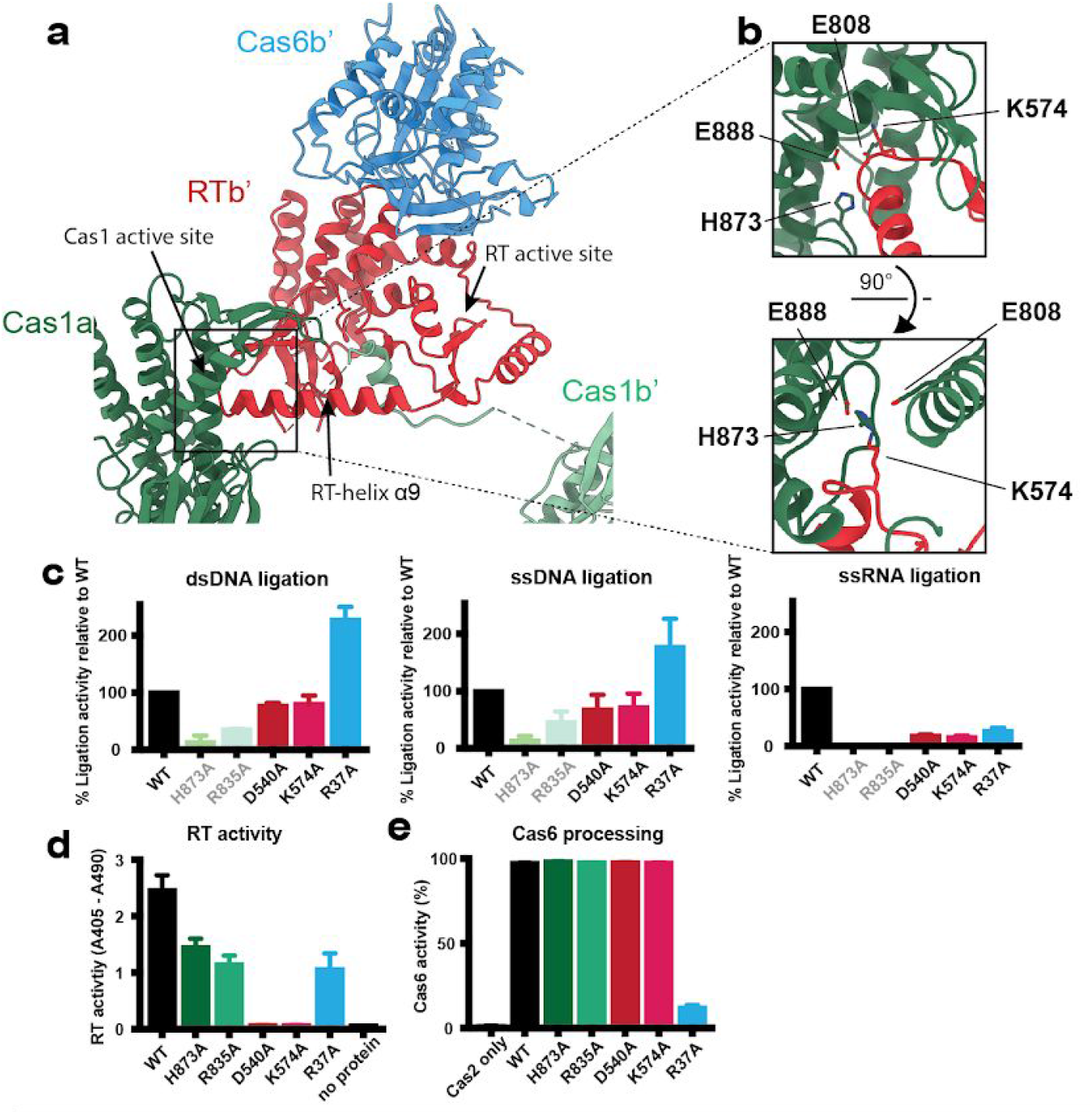
Crosstalk between RT and Cas1 and Cas6 active sites. **a)** Interaction between RT-helix α9 and Cas1 active site (color-coding from Fig. 1). **b)** Closeup of RT-helix in Cas1a and 90° rotation. Cas1 active site residues and RT-helix K574 are shown in stick configuration. **c)** Ligation of fluorescent 35-nt dsDNA (0.5 μM), ssDNA (0.5 μM), and ssRNA (4 μM) protospacers into target pCRISPR by mutant Cas6-RT-Cas1—Cas2s. The percent ligation activity is calculated as the fraction of fluorescent products from the mutant complex relative to that from the WT complex (mean ± sd, *n* = 3 independent experiments). Results are represented by colored bars: WT, black; Cas1 mutants, greens; RT mutants, reds; Cas6 mutant, blue. Cas1 mutant data is the same data shown in Figure 2e and are depicted in pale greens with gray labels. Representative gels are shown in Supplementary Fig. 7c. **d)** RT activity assays comparing the WT and mutant Cas6-RT-Cas1—Cas2s (mean ± sd, *n* = 3 independent experiments; Catalog No. 11468120910, Roche Diagnostics, Indianapolis, IN). Same color-coding is used from **c** except Cas1 mutant data is shown in darker greens with black labels. **e)** Cas6 activity assays comparing the WT and mutant Cas6-RT-Cas1—Cas2 proteins. The percent cleavage is calculated as the fraction of the fluorescent CRISPR repeat RNA that has been cleaved, with color-coding from **d** (mean ± sd, *n* = 3 independent experiments). A representative gel is shown in Supplementary Fig. 7e. Source data for panels **c-e** are available in Supplementary Data Set 3.

To investigate the functional implications of the RT-helix in the crosstalk between the RT and Cas1 domains, we generated a panel of Cas6-RT-Cas1 mutants and tested them in protospacer ligation and RT activity assays. In addition to H873 and R835, we also mutated the RT active site residue D540 and the RT-helix lysine (K574), which points into the Cas1a active site. Here, ligation activity is shown as the percent ligation into the target plasmid by the mutant Cas6-RT-Cas1—Cas2 complex relative to a parallel WT control (Fig. 6c, Supplementary Fig. 7c). Interestingly, while the D540A and K574A mutants result in a 20-30% reduction in DNA ligation, they reduce RNA ligation by more than 80%. It appears that impairing either RT activity or the RT-helix-Cas1a interaction critically disrupts RNA ligation but not DNA ligation.

The results from the RT activity assays show further evidence of crosstalk between the RT and Cas1 domains. Reverse transcriptase activity levels for the different mutant Cas6-RT-Cas1—Cas2 complexes showed that mutating either D540 or K574 abolishes RT activity. The Cas1 domain mutations also show significant effects: the H873A mutant and R835A result in a 41% and 53% reduction in dNTP incorporation, respectively, after two hours (Fig. 6d). These results show that despite being far away from the RT active site, the RT-helix lysine as well as the Cas1 residues potentially involved in the RT-helix—Cas1 interaction are significant for RT activity, supporting the hypothesis that the RT-helix plays an essential role in regulating RT activity. Together, these results suggest that there is bidirectional crosstalk between the Cas1 and RT domains.

We next explored the potential crosstalk between the Cas1/RT domains and the Cas6 domain, which lies on the opposite side of the RT. The results show that the Cas6 active site mutant R37A reduces RT activity by 57% compared to the WT (Fig. 6d). Interestingly, the R37A mutant increases DNA ligation by more than 2-fold while reducing RNA ligation by 74% compared to the WT (Fig. 6c). Though the increase in DNA ligation is puzzling, the Cas6 active site mutant may reduce RNA ligation by way of its effect on RT activity, which is more critical for RNA ligation than for DNA ligation. These results show that Cas6 is not just operating on its own and influences the other domains in the complex.

We finally tested the mutants alongside the WT Cas6-RT-Cas1 in Cas6 processing activity assays. The Cas6-RT-Cas1—Cas2 complex was incubated with a 5’-labeled 35-nt RNA substrate corresponding to the CRISPR repeat sequence. The results show that the WT Cas6-RT-Cas1—Cas2 processes the 35-nt RNA substrate down to a 26-nt product with 98% efficiency after two hours (Fig. 6e, Supplementary Fig. 7d,e). Mutating the Cas6 active site residue R37 reduces processing efficiency to only 12%. The other mutants show no difference in processing relative to the WT, suggesting that the crosstalk between the Cas6 domain and the other two domains is unidirectional, with the Cas6 active site mutation significantly affecting ligation and RT activity but the Cas1 and RT domain mutations having no effect on Cas6 RNA processing activity.

## DISCUSSION

RT-Cas1 and associated Cas2 proteins represent a unique CRISPR adaptation module shown to be involved in the acquisition of CRISPR spacers directly from RNA in a fascinating example of information flow from RNA to DNA^20,22–29,49^. In this study, we identify a type III *Thiomicrospira* Cas6-RT-Cas1—Cas2 complex containing three active sites for CRISPR spacer integration, reverse transcription, and RNA processing, and we observe all three activities in our reconstituted system. The structure of this Cas6-RT-Cas1—Cas2 complex revealed interesting differences between its Cas1—Cas2 integrase core and that observed for the *E. coli* Cas1—Cas2 integrase. An RT-helix links the RT and Cas1 active sites but also appears to block the position of substrate binding. Functional interactions between the Cas6, RT, and Cas1 domains show bidirectional crosstalk between the RT and Cas1 domains and unidirectional crosstalk from Cas6 to the other two domains.

Although the *Thiomicrospira* Cas6-RT-Cas1—Cas2 complex catalyzes site-specific cleavage-ligation chemistry with dsDNA, ssDNA, and ssRNA substrates, we only observe full-site integration with dsDNA substrates. Its inability to fully integrate single-stranded substrates even in the presence of dNTPs implies that additional host factors may be required to carry out this process *in vivo*. It is still unclear when the RNA substrate is reverse transcribed into cDNA during spacer acquisition. We found that the RT faithfully synthesizes cDNA from primer/template hybrids with short ~35-nt RNA or DNA templates and with minimal homology requirements; however, the preferred template and primer for these systems *in vivo* remain unknown. Though previous work on the MMB-1 system suggested that the cleavage-ligation products could be substrates for target-primed reverse transcription similar to group II intron retrohoming^27,49,50^, we did not see evidence of this in biochemical assays. If such a process were to occur *in vivo,* it is interesting to speculate whether the “twisted” arrangement of the Cas1—Cas2 components relative to their observed positioning in the *E. coli* Cas1—Cas2 integrase might be advantageous to stall integration at some intermediate step to allow access for the RT.

The structural and functional interactions detected between the Cas1, RT, and Cas6 active sites suggest a model in which the linking RT-helix serves as a regulator of both RT and Cas1 activities through its movement in and out of the Cas1 active site (Fig. 7a,b). Given the RT-helix connection to the β-sheet containing the RT active site, we speculate that the motion of the RT-helix drives the RT to adopt a conformation more closely resembling the active conformation of other RT structures, or vice versa, that the conformational change of the β-sheet drives the motion of the RT-helix (Fig. 7a). We suggest that the Cas6b-RTb and Cas6b’-RTb’ domains may be able to adopt two states: one where the Cas6-RT lobe is freely moving, only flexibly attached by a linker to the Cas1 domain in the same chain and another where it docks onto the opposite Cas1a domain, blocking Cas1 from carrying out integration (Fig. 7b). Having on-off switches for the RT and Cas1 active sites may be critical to the process of RNA spacer acquisition where intermediate substrates may pass from one active site to the other. Additionally, the unidirectional crosstalk between the Cas6 to the other two domains also implies a possible multi-step process in which coordination between the different enzymatic functions creates an ordered process for spacer integration and CRISPR RNA processing.

**Figure 7.**
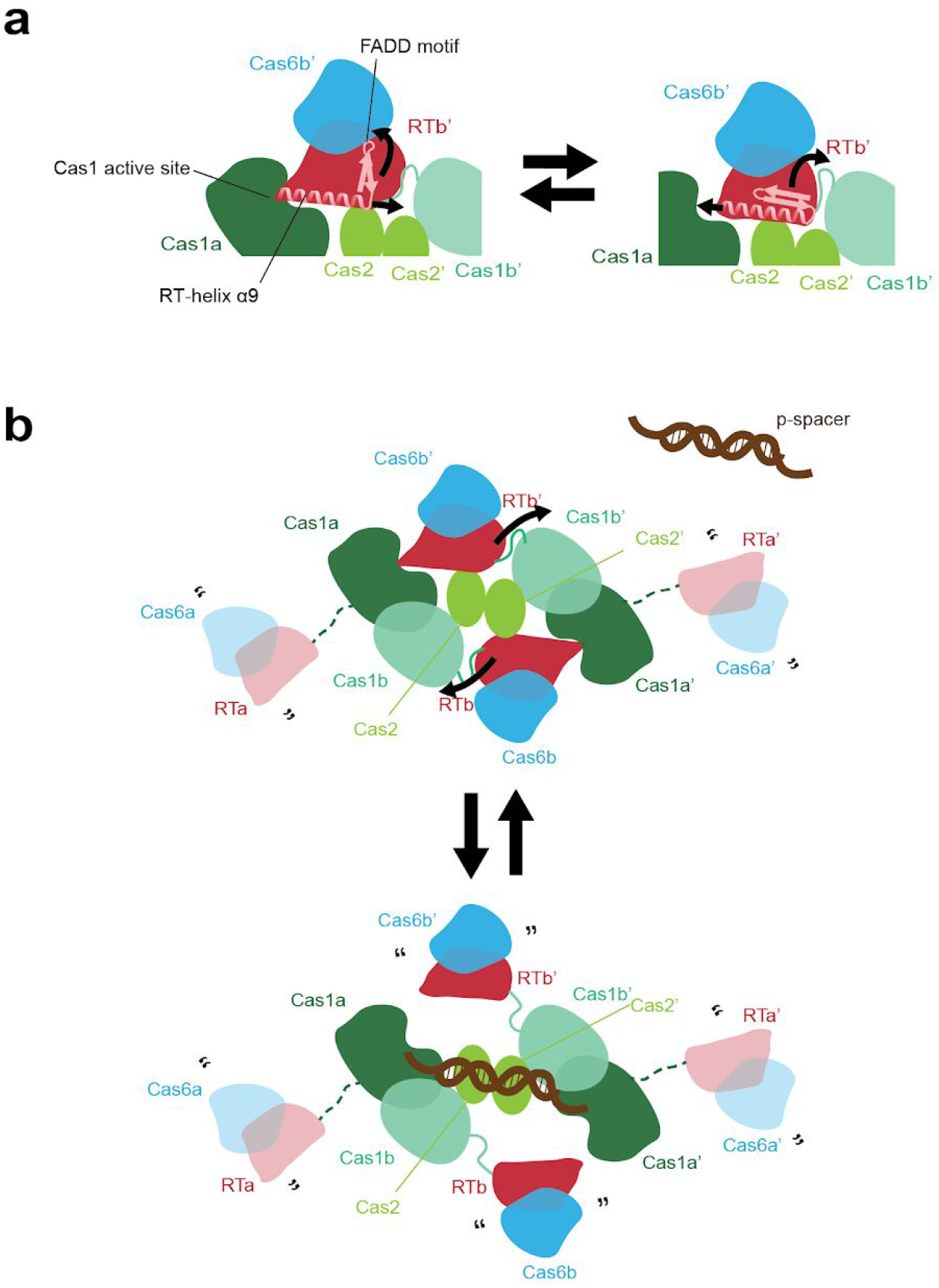
Model for RT activation. **a)** Cartoon showing hypothesized motion of RT-helix α9 in and out of the Cas1a active site and the hypothesized motion of the β-strands attached to the FADD motif containing the active site residues. Proteins colored by domain: Cas6, blue; RT, red, Cas1a, dark green; Cas1b, turquoise; Cas2, lime green. Visible RT-Cas1 linkers are shown as solid turquoise lines. RT-helix α9 and connecting β-strands containing FADD motif are schematized and shown in pink. Hypothesized motions are indicated by curved arrows. **b)** Cartoon showing hypothesized conformational change of the Cas6-RT-Cas1—Cas2 complex to accommodate a protospacer substrate, with color-coding from **a**. Missing Cas6-RT domains (Cas6a/RTa and Cas6a’/RTa’) that are present in the complex but not visible are depicted as semi-transparent, connected to the rest of the structure by dashed dark green lines. Quotation marks “” indicate presumed mobility. Protospacer substrate is schematized in brown. Hypothesized motions are indicated by curved arrows.

Our results imply that Cas6-RT-Cas1—Cas2 functions as a carefully regulated system to enable spacer acquisition from DNA and potentially RNA sources. Questions remain about the relative efficiency of these reactions in light of our observations that the complex strongly prefers DNA over RNA substrates for cleavage-ligation, suggesting that RNA may not be the primary source of spacers for this system. Previous literature examining correlation between the frequency of spacer acquisition and overall gene expression level does indicate that there is variation among these systems *in vivo*^27-29^. It is also possible that the efficiency in integrating RNA substrates varies depending on the substrate as a part of a selection process to distinguish between self and non-self. Future research on RNA spacer acquisition by these RT-Cas1 and Cas6-RT-Cas1 fusions can more broadly shed light on how these systems may potentially generate immunity against RNA invaders, which is still a topic that is not very well understood. Furthermore, deeper insight into these fusions can help make strides toward advancing tools that utilize these proteins to record transcriptional information in cells. As the first high-resolution structure of the Cas6-RT-Cas1—Cas2 complex, our structure provides key details toward achieving a greater understanding of RT-associated integrases and their functions.

## METHODS

### Cas6-RT-Cas1 Protein Identification

Hidden Markov Models were built from RT-Cas1 fusion proteins or Cas1 sequences affiliated with RTs obtained from a previous study^26^ and used to search for more RT-Cas1 representatives. Metagenomic contigs harboring fusion proteins consisting of RT, Cas1, and Cas6 domains were selected from a groundwater, CO_2_-driven Geyser dataset and curated for sequencing errors as previously described^54^, and a Cas6-RT-Cas1 protein and its cognate CRISPR array and Cas2 from a *Thiomicrospira* genome assembly became the subject of this study. The CRISPR locus is available in Supplementary Data Set 1.

### Residue Conservation Analysis

A diverse set of 968 Cas1 proteins and 92 Cas1s fused to or associated with RTs (and in some cases, Cas6) was used for analysis of the conservation of the cluster of residues consisting of R832, R834, R835, and H879. An alignment was generated from the aforementioned set with the Cas6-RT-Cas1 protein of interest using the MAFFT v7.407 LINSI setting with 1000 iterations^55^. The positions of interest were extracted and a sequence logo was generated to display the conservation of residues across the relevant Cas1 proteins.

### Protein Purification

The Cas6-RT-Cas1 and Cas2 genes were codon optimized for *E. coli* expression and ordered as G-blocks. Cas6-RT-Cas1 and Cas2 were PCR amplified and cloned separately into a pET-based expression vector with an N-terminal 10xHis-MBP-TEV tag. Plasmids were transformed into chemically competent Rosetta cells. Cells were grown to an OD_600_ of ~0.6 and induced overnight at 16 °C with 0.5 mM isopropyl-β-D-thiogalactopyranoside (IPTG). Cells were harvested and resuspended in lysis buffer (20 mM HEPES, pH 7.5, 500 mM NaCl, 10 mM imidazole, 0.1% Triton X-100, 1 mM Tris (2-carboxyethyl)phosphine (TCEP), Complete EDTA-free protease inhibitor (Roche), 0.5 mM phenylmethylsulfonyl fluoride (PMSF), and 10% glycerol. After cell lysis by sonication, lysate was clarified by centrifugation and the supernatant was incubated on Ni-NTA resin (Qiagen). The resin was washed with wash buffer (20 mM HEPES, pH 7.5, 500 mM NaCl, 10 mM imidazole, 1 mM TCEP, and 5% glycerol), before the protein was eluted with wash buffer supplemented with 300 mM imidazole and then digested with TEV protease overnight. The salt concentration was diluted to 335 mM NaCl using ion-exchange buffer A (20 mM HEPES, pH 7.5, 1 mM TCEP, and 5% glycerol), the cleaved MBP tag was removed with an MBPTrap column (GE Healthcare), and the protein was bound to a HiTrap heparin HP column (GE Healthcare), before elution with a gradient from 335 mM to 1 M KCl. The protein was then concentrated and purified on a Superdex 200 (16/60) column with storage buffer (20 mM HEPES, pH 7.5, 500 mM KCl, 1 mM TCEP, and 5% glycerol). The same purification protocol was used for Cas6-RT-Cas1 (wild-type and mutants) and Cas2. Sequences of proteins are shown in Supplementary Table 1.

### DNA and RNA Substrate Preparation

To generate target plasmid pCRISPR, the CRISPR array was reduced and ordered as two fragments, which were amplified by PCR and inserted into a pUC19 backbone by Gibson Assembly. For the full-site selection, pCRISPR_full-site was generated by cloning a *cat* promoter and chloramphenicol resistance gene into a pUC19 plasmid, removing the RBS sequence and start codon, and then inserting a sequence carrying 163 bp of the leader from pCRISPR and the 35 bp repeat before the chloramphenicol resistance gene. DNA and RNA oligos used in this study were ordered from Integrated DNA Technologies (IDT) and purified on 6% (for >35-nt oligos) or 14% urea-PAGE (for ≤35-nt oligos). Protospacers, dsDNA targets, primer/template hybrids, and the half-site substrate were formed by heating at 95 °C for 5 minutes and slow cooling to room temperature in either HEPES hybridization buffer (20 mM HEPES, pH 7.5, 25 mM KCl, and 10 mM MgCl_2_) or Tris hybridization buffer (20 mM Tris, pH 7.5, 25 mM KCl, and 10 mM MgCl_2_), as consistent with the integration buffer in which the substrate will be used. For the half-site substrate, hybridization was carried out with a 1.25-fold excess of the shortest strand before purification on 8% native PAGE. Sequences of the substrates are shown in Supplementary Table 2.

### Integration Assays

Integration assays with supercoiled target plasmid were conducted in HEPES integration buffer (20 mM HEPES, pH 7.5, 25 mM KCl, 10 mM MgCl_2_, 1 mM DTT, 0.01% Nonidet P-40, and 10% DMSO), while integration assays with the short linear dsDNA target were conducted in Tris integration buffer (20 mM Tris, pH 7.5, 25 mM KCl, 10 mM MgCl_2_, 1 mM DTT, 0.01% Nonidet P-40, and 10% DMSO). Cas6-RT-Cas1 and Cas2 were first pre-complexed at 2 μM at room temperature for 30 min. in the designated integration buffer. The protospacer substrate was then incubated with the Cas6-RT-Cas1 and Cas2 complex for 15 minutes, followed by addition of target plasmid (pCRISPR or pCRISPR_full-site) to 20 ng/mL (~10 nM) or short linear dsDNA target (250 nM). Integration reactions with ssRNA protospacers are supplied with 1 mM dNTPs unless otherwise indicated. The reaction was carried out at 37 °C for 2 hours. A 4 μM and 1 μM concentration was used for all protospacers for the time-course integration assays and integration assays with a short dsDNA linear target, respectively. For all other integration assays, a 500 nM concentration is used for dsDNA and ssDNA protospacers and a 4 μM concentration is used for ssRNA protospacers. In reactions where fluorescent protospacers are indicated, oligos with a 5’ 6-carboxyfluorescein (FAM) attachment were ordered. The fluorescent dsDNA protospacers were generated by hybridizing one 5’-labeled strand with its complementary unlabeled strand. For reactions with a supercoiled target plasmid, the reaction was quenched by the addition of 0.4% SDS and 25 mM EDTA, treated with proteinase K for 15 minutes at room temperature, and then treated with 3.4% SDS, before analysis on a 1.5% agarose gel (unstained for reactions with fluorescent protospacers and pre-stained with SYBR Safe for reactions with unlabeled protospacers). For reactions with a short linear dsDNA target, the reaction was quenched by the addition of 2 vol. of quench buffer (95% formamide, 30 mM EDTA, 0.2% SDS, and 400 μg/mL heparin) and heating at 95 °C for 4 min., before analysis on a 6% urea-PAGE gel. The same amount of reaction product is added to each lane for all gel analyses. When examining integration by open-circle formation, gel bands were visualized by ChemiDoc MP (BioRad) and quantified using Image Lab (BioRad). The fraction plasmid nicked was calculated as the ratio of the open-circle integration product band intensity to the total intensity of both the open-circle band and the supercoiled plasmid band. Fluorescent bands were visualized by Typhoon FLA gel imaging scanner and the band intensities are quantified using ImageQuantTL. The relative percent ligation activity of the mutant proteins was calculated as the ratio of the product band intensity from the mutant protein complex relative to the product band intensity from the parallel WT control. Statistical analyses were performed using Prism GraphPad. Data were presented as the mean ± SD (error bars) of three independent experiments.

For full-site integration assays, pCRISPR_full-site is used as the target plasmid. Sixteen 50 uL integration reactions were carried out as described above with the reporter construct for 2 hours. Instead of quenching the reaction, the integration products were purified using the Qiagen MinElute PCR Purification Kit and electroporated into DH10B cells. Transformants were plated on LB agar containing chloramphenicol and 95 of the surviving colonies were sequenced using Sanger Sequencing.

### RT and Cas6 Activity Assays

RT activity was measured using an ELISA-based colorimetric reverse transcriptase activity assay (Catalog No. 11468120910, Roche Diagnostics, Indianapolis, IN). The HEPES integration buffer was used for the reaction. Cas6-RT-Cas1 and Cas2 were first pre-complexed at 2 μM at room temperature for 30 min. The reactions were conducted in triplicates using the supplied template/primer hybrid poly (A) × oligo (dT)_15_ at 37 °C for 2 hours. After following the assay kit’s ELISA procedure, the RT activity was measured in absorption units (A_405 nm_ - A_490 nm_) using a fluorescence plate reader (BioTek). Statistical analyses were performed using Prism GraphPad. Data were presented as the mean ± SD (error bars) of three independent experiments.

For template-driven cDNA synthesis reactions off a fluorescent DNA primer annealed to a DNA or RNA template, Cas6-RT-Cas1 and Cas2 were first pre-complexed at 2 μM at room temperature for 30 min before incubating with 250 nM primer/template substrate and 1 mM dNTPs, where indicated, at 37 °C for 2 hours. For Cas6 activity assays, Cas6-RT-Cas1 and Cas2 were first pre-complexed at 2 μM at room temperature for 30 min. before incubating with the 5’-labeled 35-nt RNA substrate corresponding to the repeat sequence at 37 °C for 2 hours. The template-driven cDNA synthesis reactions and Cas6 processing reactions were quenched by adding 2 vol. quench buffer and heating at 95 °C for 4 min., before analysis on a 14% urea-PAGE gel. Fluorescent bands were visualized by Typhoon FLA gel imaging scanner. The percent Cas6 activity is quantified as the ratio of the product band intensity to the total intensity of both the product band and the unprocessed RNA substrate band.

### Grid Preparation

Cas6-RT-Cas1—Cas2 and the half-site DNA substrate were complexed by mixing 50 μM Cas6-RT-Cas1, 50 μM Cas2, and 12.5 μM DNA half-site substrates in storage buffer and dialyzing in complex buffer (10 mM HEPES, pH 7.5, 5 mM EDTA, 250 mM KCl, and 1 mM TCEP) for 2 hours using the Slide-A-Lyzer MINI Dialysis Devices and concentrated to 100 μM Cas6-RT-Cas1—Cas2. For freezing grids, a 3 μl drop of protein was applied to freshly glow discharged Holey Carbon, 300 mesh R 1.2/1.3 gold grids (Quantifoil, Großlöbichau, Germany). A FEI Vitrobot Mark IV (ThermoFisher Scientific) was used with 4°C, 100% humidity, 1 blot force, and a 3 second blot time. Grids were then clipped and used for data collection.

### Cryo-EM Data Acquisition

Grids were clipped and transferred to a FEI Titan Krios electron microscope operated at 300 kV. 50 frame movies were recorded on a Gatan K3 Summit direct electron detector in super-resolution counting mode with pixel size of 0.5953 Å. The electron dose was 10.50 e^-^ Å^2^ s^-1^ and total dose was 49.98 e^-^ Å^2^. Initial data collection was performed with a GIF Quantum energy filter inserted, however after 516 micrographs had been collected it became unstable and was removed for the final 2729 micrographs. Nine movies were collected around a central hole position with image shift and defocus was varied from −1 to −3 μm through SerialEM^56^. See Supplementary Table 3 for data collection statistics.

### Cryo-EM Data Processing

Motion-correction and dose-weighting were performed on all 3330 movies 2x “binned” to 1.187 Å per pixel using RELION 3.0’s implementation of MotionCor2^57,58^. CTFFIND-4.1 was used to estimate the contrast transfer function (CTF) parameters^59^. Micrographs were then manually examined to remove subjectively bad micrographs, such as empty or contaminated holes, resulting in 2,035 micrographs^60^. Additionally, micrographs with a CTF maximum resolution lower than 4 Å were discarded, yielding 1998 micrographs. Template-free auto-picking of particles was performed with RELION3.1’s Laplacian-of-Gaussian (LoG) filter to generate an initial set of particles which were iteratively classified to find a subset of particles that were subsequently used to template-based auto-pick 249,102 particles.

Template picked particles were iteratively 2D-classified in RELION3.0 resulting in a set of 129,175 particles. These particles were imported to cryoSPARC v2. An initial map was generated with an ab-initio job and refined with subsequent homogeneous and non-uniform refinement jobs^61,62^. ‘csparc2star.py’ from UCSF pyem was used to convert the resulting data to a RELION star file^63^. These particles were then refined in RELION3.0 resulting in a map with ~4.1 Å overall resolution. Subsequent refinement following Bayesian particle polishing, and CTF refinement resulted in a map with ~3.66 Å overall resolution^64^. A mask surrounding the stable side of the complex was generated and supplied to a refinement resulting in a map with ~3.42 Å overall resolution. Further classification in 3D was attempted but did not result in an improved map, nor did the application of C2 symmetry during refinement.

This set of 129,175 particle coordinates were used for training in the Topaz particle-picking pipeline^65^. Training, picking, and extraction in Topaz resulted in 1,305,497 particles. These particles were then processed using a similar pipeline to the above Relion template-picked particles, resulting in 285,342 particles after 2D classification and yielding a ~3.8 Å map before, and a ~3.3 Å map following Bayesian particle polishing. The final maps were not distinguishably improved compared to the template-picked particle set. As with the prior particle set, further classification in 3D could not identify a subset of particles yielding an improved map. A consensus homogeneous refinement of these particles in cryoSPARCv2 was used as input for 3D variability analysis in cryoSPARCv2 (filter resolution 5 Å)^66^.

### Modeling, Refinement, and Analysis

De novo modeling of the best-resolved portion of the complex was performed in Coot using the output map from a masked 3D refinement in Relion sharpened with Phenix.auto_sharpen^67,68^. The *de novo* modeled structure, corresponding to Cas2 chain A (amino acids 1-90), Cas2 chain B (amino acids 1-96), Cas6-RT-Cas1 chain C (amino acids 1-634), Cas6-RT-Cas1 chain D (amino acids 650-981), and Cas6-RT-Cas1 chain E (amino acids 641-981), was refined using Phenix.real_space_refine. Molprobity was used to guide iterative rounds of manual adjustment in Coot and refinement in Phenix^69^. Subsequently, portions of the refined model were copied and rigid-body docked into density corresponding to less-well resolved portions of the complex (corresponding to Cas2 chain A (amino acids 91-96), Cas6-RT-Cas1 chain C (amino acids 641-981), Cas6-RT-Cas1 chain E (amino acids 1-634), and Cas6-RT-Cas1 chain F (amino acids 650-981)) in a postprocessed Relion 3D refinement output map that was masked to include the entire visible complex. A final round of rigid-body refinement was performed in Phenix.real_space_refine. This model consists of two entire Cas2 protomers (amino acids 1-96), two entire Cas6-RT-Cas1 protomers (amino acids 1-981), and two partial Cas6-RT-Cas1 protomers with only the Cas1 domain modeled (amino acids 650-981). Six loops were unmodeled in Cas6-RT-Cas1 protomers: three in the Cas6 domain (amino acids 95-110, 248-257, and 212-221), two in the RT domain (amino acids 383-388 and 593-616), and a linker between the RT and Cas1 domains (amino acids 635-640). Figures were prepared using Chimera, ChimeraX, Prism, GNU Image Manipulation Program, and Adobe Illustrator software^70,71^.

### Data and reagent availability

The atomic model of the partial Cas6-RT-Cas1—Cas2 complex masked for the stable density is in the Protein Data Bank (PDB) under 7KFT and the corresponding map is deposited in the Electron Microscopy Data Bank (EMDB) under EMD-22855. The atomic model of the full Cas6-RT-Cas1—Cas2 complex is in the Protein Data Bank (PDB) under 7KFU and the corresponding map is deposited in the Electron Microscopy Data Bank (EMDB) under EMD-22856. The original micrograph movies and final particle stack are deposited in the Electron Microscopy Public Image Archive (EMPIAR) under EMPIAR-XXXXX. The uncropped images for the main text or supplementary figures are available in Supplementary Data Set 2. Source data for Fig. 2c,f, 3c,d, 4c, 5d, 6c-e are available in Supplementary Data Set 3. Source data for all other experiments that support the findings of this study are available from the corresponding author upon reasonable request.

## Supporting information

Supplementary Data Set 3

Supplementary Data Set 2

Supplementary Data Set 1

## ACKNOWLEDGMENTS

We thank Paul Tobias for computational resources at the Cal-Cryo EM facility, and Dr. James Hurley and Dr. Eva Nogales for supporting the microscopy work. This material is based upon work supported by the National Science Foundation under award number 1817593. J.Y.W. is supported by the US National Science Foundation Graduate Fellowship and previously by the Berkeley Graduate Fellowship. B.A.-S. is supported by the US National Science Foundation Graduate Fellowship. S.G.B. is a New York Stem Cell Foundation Robertson Neuroscience Investigator. J.A.D. is an investigator of the Howard Hughes Medical Institute. We thank A.V. Wright for input on the manuscript and members of the Doudna laboratory and Brohawn laboratory for comments and discussions.

## AUTHOR CONTRIBUTIONS

J.Y.W. conceived of and designed experiments with input from C.M.H., J.A.D., and S.G.B. J.Y.W. performed protein expression, purification, and biochemical experiments. C.M.H. and J.Y.W. performed cryo-EM data collection, and C.M.H. and S.G.B. processed the cryo-EM data and built and refined the atomic models. B.A.-S. identified the CRISPR-Cas locus of interest and performed bioinformatics experiments and analyses with input from J.F.B. J.Y.W., C.M.H., and J.A.D. wrote the manuscript and all authors edited the manuscript.

## COMPETING INTERESTS

The Regents of the University of California have patents issued and pending for CRISPR technologies on which J.A.D. is an inventor. J.A.D. is a cofounder of Caribou Biosciences, Editas Medicine, Scribe Therapeutics, Intellia Therapeutics and Mammoth Biosciences. J.A.D. is a scientific advisory board member of Caribou Biosciences, Intellia Therapeutics, eFFECTOR Therapeutics, Scribe Therapeutics, Mammoth Biosciences, Synthego, Algen Biotechnologies, Felix Biosciences and Inari. J.A.D. is a Director at Johnson & Johnson and has research projects sponsored by Biogen, Pfizer, AppleTree Partners and Roche.

## Supplementary Information

**Supplementary Figure 1.**
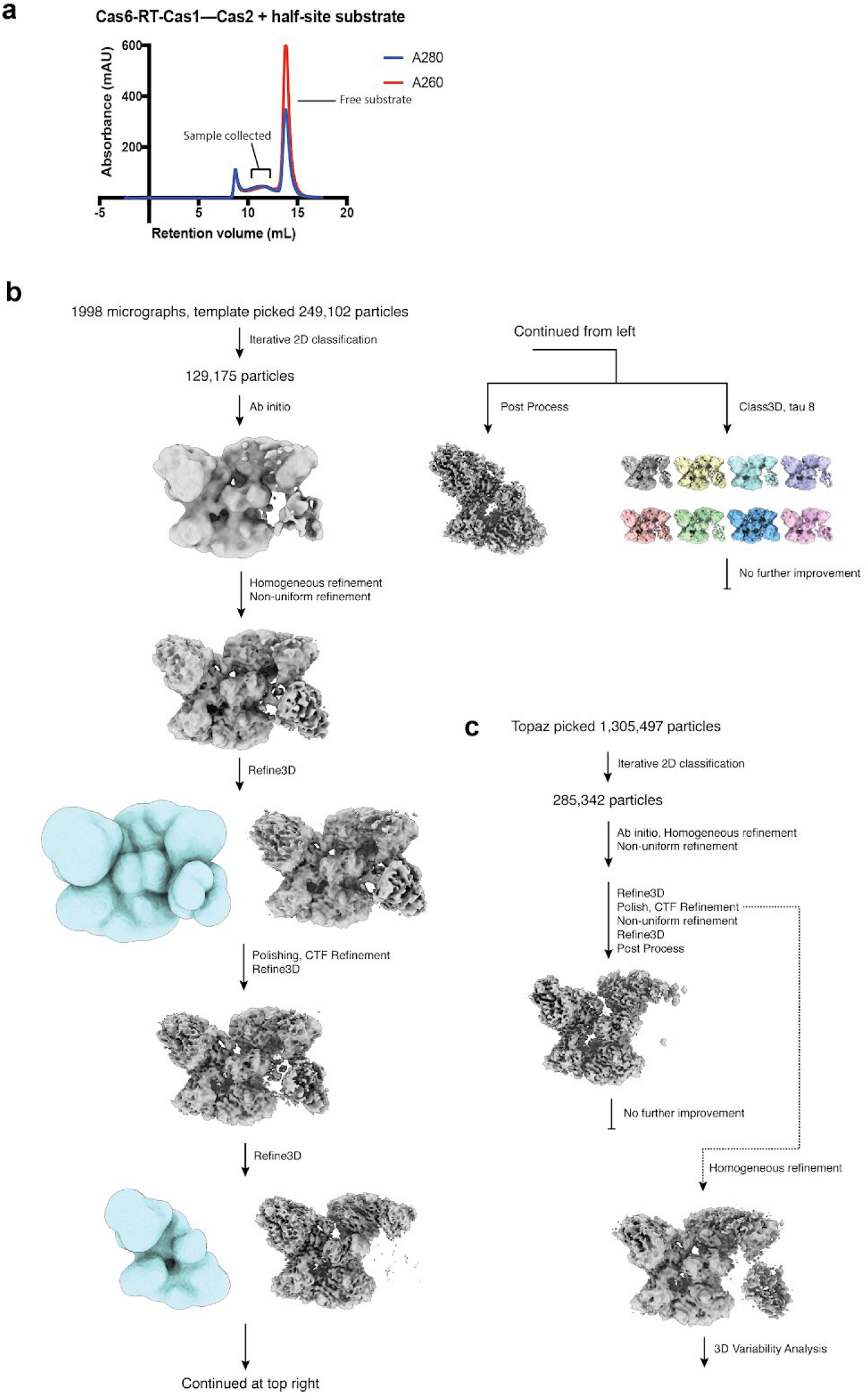
Complex formation and Cryo-EM processing pipeline. **a)** Gel filtration run of Cas6-RT-Cas1—Cas2 complexed to DNA half-site substrate. The peaks representing the sample used for cryo-EM data collection and free substrate are indicated. **b,c)** Cryo-EM data processing pipeline in RELION and cryoSPARC from RELION picked particles and Topaz picked particles respectively. Cryo-EM maps are colored gray, and masks used in RELION colored light blue. See Methods for details.

**Supplementary Figure 2.**
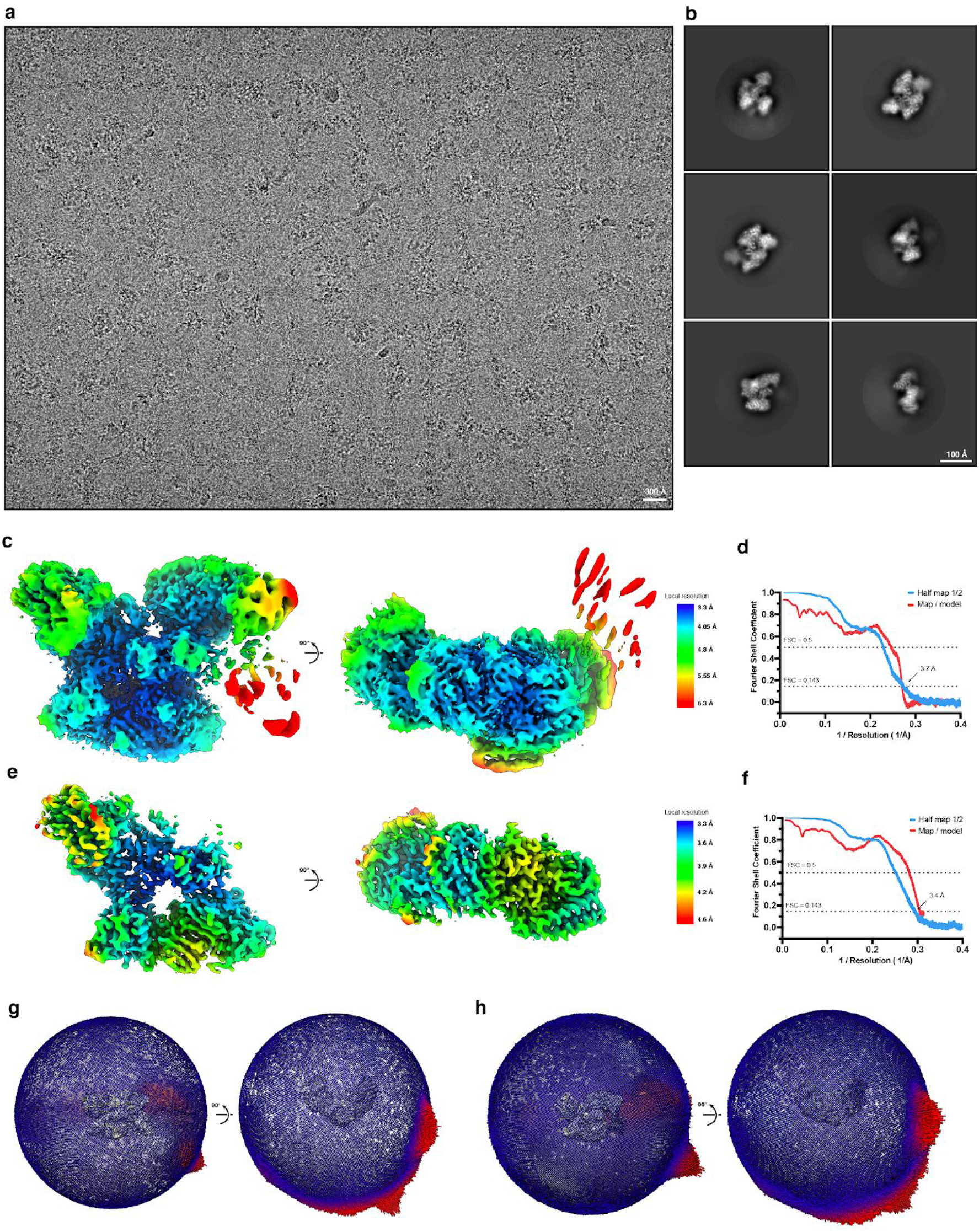
Cryo-EM validation. **a)** Representative micrograph with 10 Å lowpass filter. **b)** Selected 2D class averages. **c,e)** Local resolution estimated in RELION3.1 with map surfaces colored in rainbow as indicated, for full and partial complexes respectively. **d,f)** Fourier Shell Correlation (FSC) for the two unfiltered half-maps (light blue) and final map versus model (red), for full and partial complexes respectively. **g,h)** Angular distribution of particles with corresponding cryo-EM map, for full and partial complexes respectively.

**Supplementary Figure 3.**
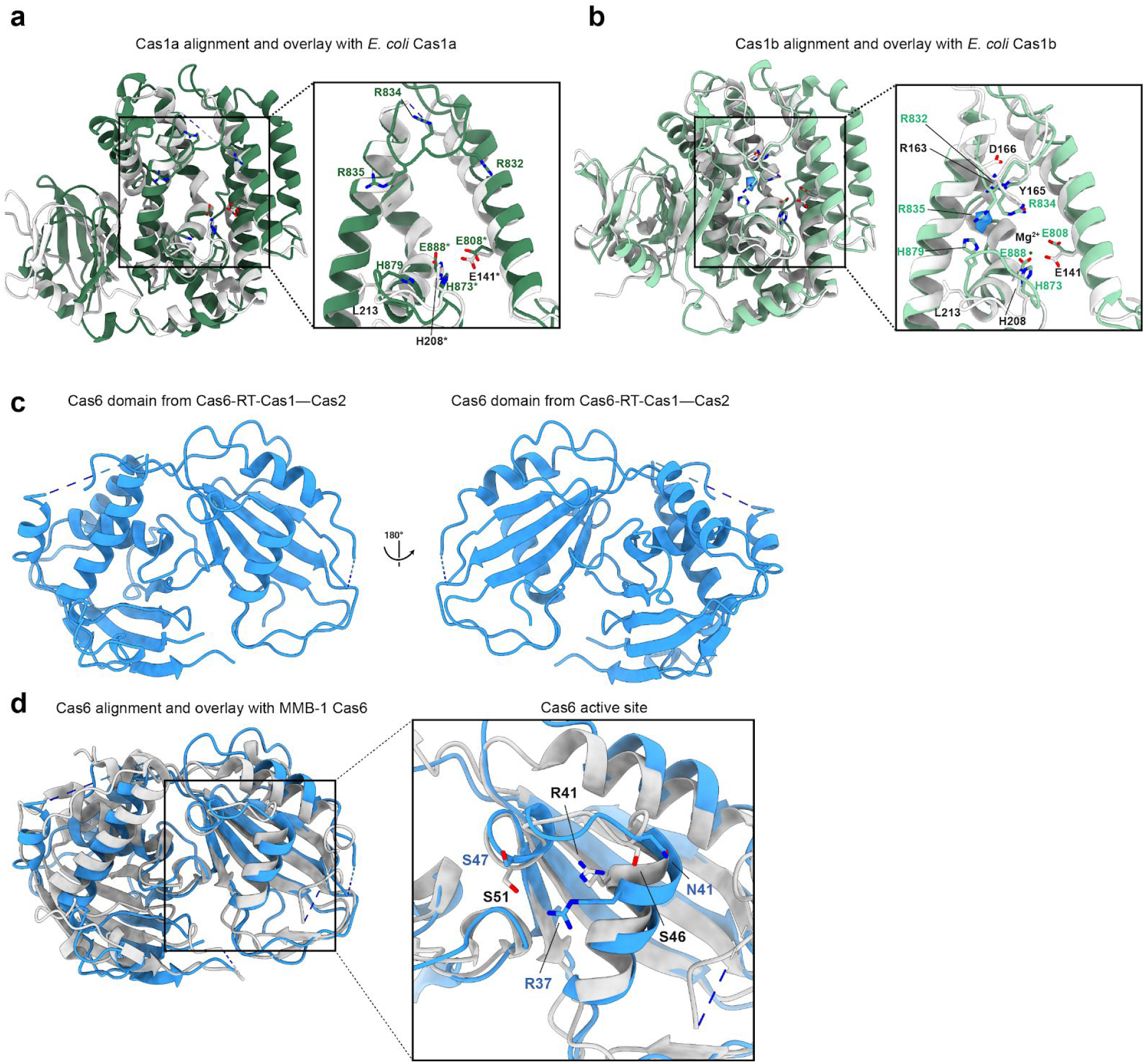
Architecture of Cas1 and Cas6 domains. **a)** Alignment and overlay of Cas1a domain (dark green) from Cas6-RT-Cas1—Cas2 structure with *E. coli* Cas1a (white, PDB: 5DS5) and closeup of active site and surrounding residues, shown in stick configuration. Cas6-RT-Cas1—Cas2 residues are labeled in green, *E. coli* Cas1 residues labeled in black. Active site residues are labeled with an asterisk *. **b)** Alignment and overlay of Cas1b domain (turquoise) with *E. coli* Cas1b (white, PDB: 5DS5) and closeup of active site and surrounding residues. **c)** Architecture of Cas6 domain from Cas6-RT-Cas1—Cas2 structure and 90° rotation. **d)** Alignment and overlay of Cas6 domain from Cas6-RT-Cas1—Cas2 structure (blue) with Cas6 domain (PDB: 6DD5) from MMB-1 Cas6-RT-Cas1 fusion (white) and closeup of active site residues, shown in stick configuration. Cas6-RT-Cas1—Cas2 residues are labeled in blue, MMB-1 Cas6 residues labeled in black.

**Supplementary Figure 4.**
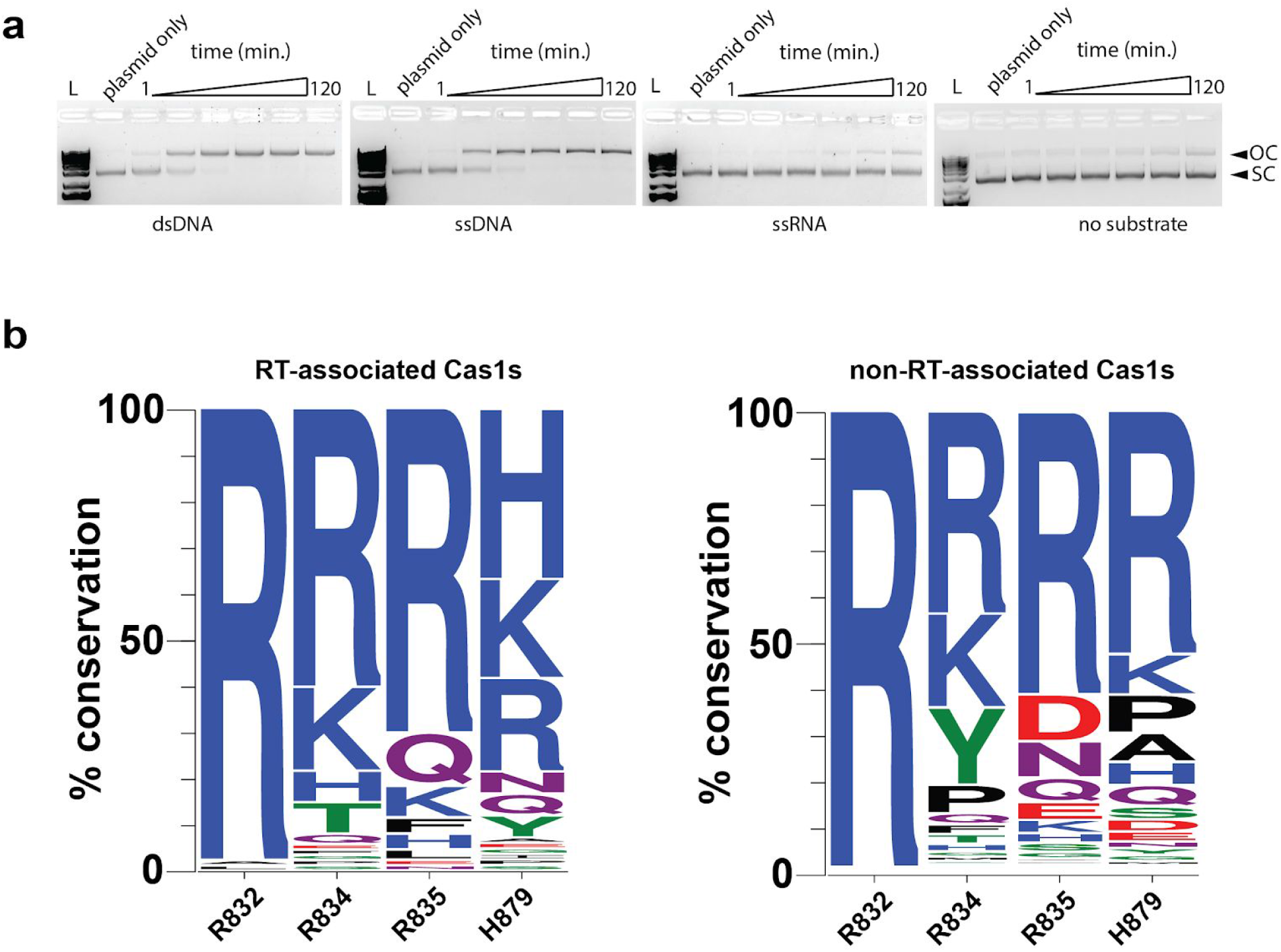
Cas6-RT-Cas1—Cas2 prefers DNA substrates over RNA substrates for cleavage-ligation. **a)** Time-course integration reactions with 35-nt dsDNA, ssDNA, ssRNA protospacers (4 M) and a control with no protospacer. The open-circle integration products (OC) and supercoiled target plasmid (SC) are indicated. **b)** Sequence logos indicating the most conserved residues of the R832, R834, R835, and H879 motif in RT-associated Cas1s (left) and non-RT-associated Cas1s (right) following sequence alignment across a diverse selection of Cas1s.

**Supplementary Figure 5.**
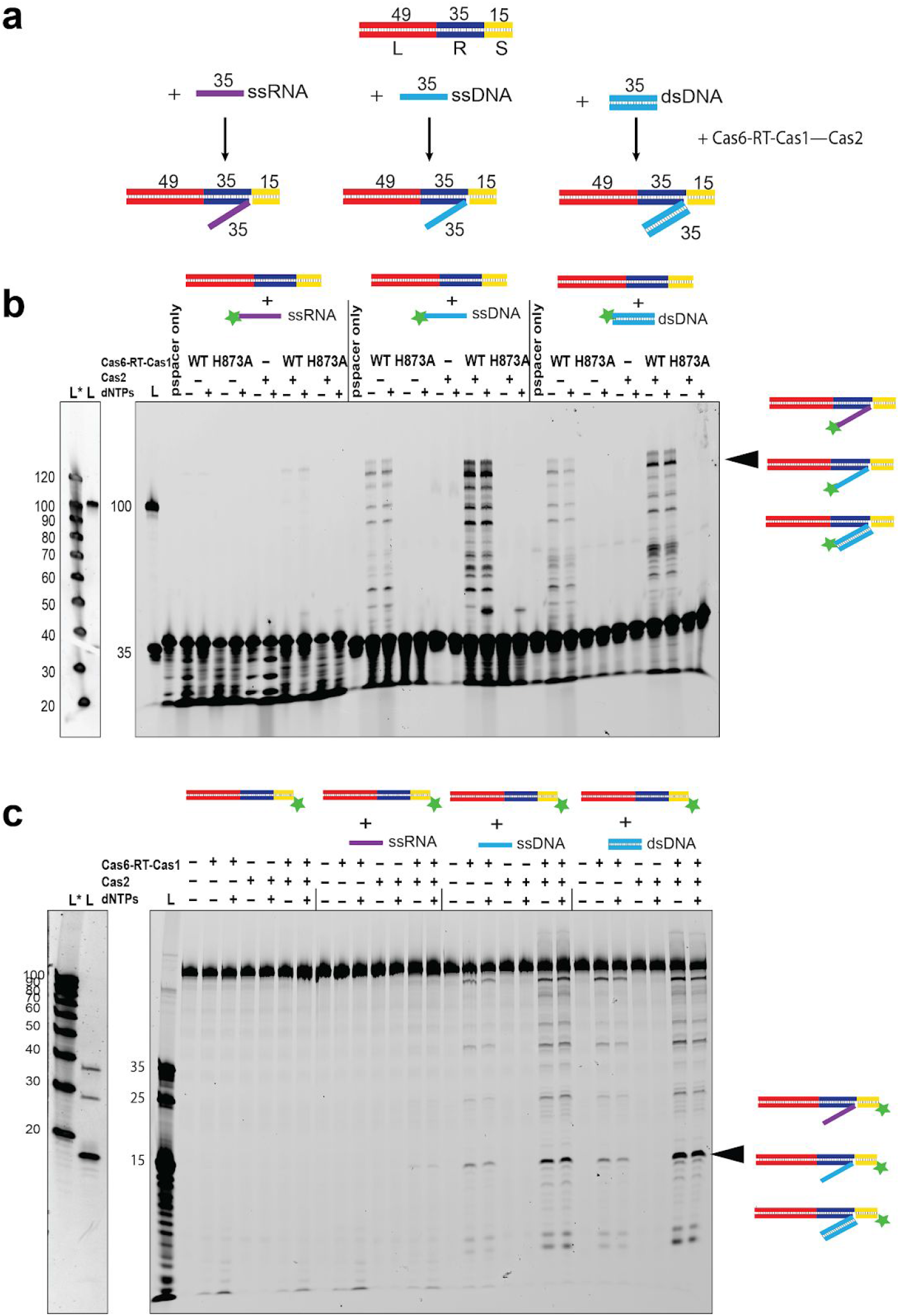
Cas6-RT-Cas1—Cas2 catalyzes ligation of DNA and RNA protospacers into the CRISPR array. **a)** Schematic of in vitro integration reaction of DNA or RNA protospacer into a short linear dsDNA target containing the CRISPR repeat. The lengths of the leader (L), repeat (R), spacer (S), and protospacers are indicated. **b)** Ligation of fluorescent dsDNA, ssDNA, and ssRNA protospacers (1 M) into a short linear target containing the CRISPR repeat. Expected products are indicated. Star indicates 6-carboxyfluorescein label. On the left, a non-fluorescent ladder L* is shown next to the fluorescent ladder L, visualized after SYBR Gold poststaining. **c)** Ligation assays conducted with fluorescent target (bottom strand labeled). Uncropped gels are available in Supplementary Data Set 2.

**Supplementary Figure 6.**
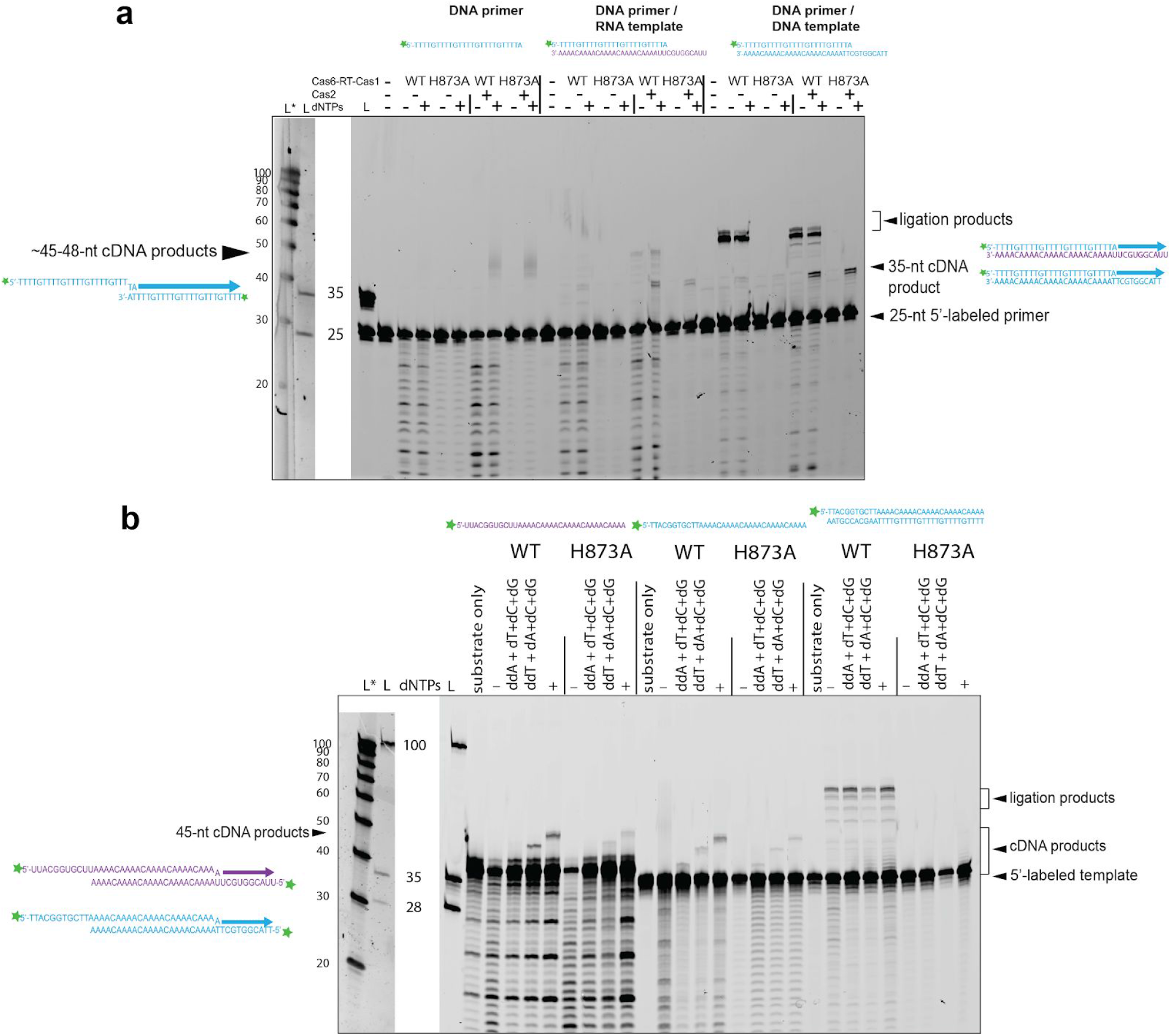
Cas6-RT-Cas1 catalyzes cDNA synthesis with minimal primer-template homology. **a)** Template-driven cDNA synthesis reactions off a fluorescent DNA primer annealed to DNA and RNA templates with WT and H873A Cas6-RT-Cas1 complexed with Cas2. Substrate sequences and expected cDNA synthesis reactions are indicated. Star indicates 6-carboxyfluorescein label. On the left, a non-fluorescent ladder L* is shown next to the fluorescent ladder L, visualized after SYBR Gold poststaining. **b)** Template-driven cDNA synthesis reactions in the absence of different dNTPs and in the presence of added ddNTPs with DNA or RNA template with no pre-annealed primer. Expected cDNA synthesis reactions are indicated. Uncropped gels are available in Supplementary Data Set 2.

**Supplementary Figure 7.**
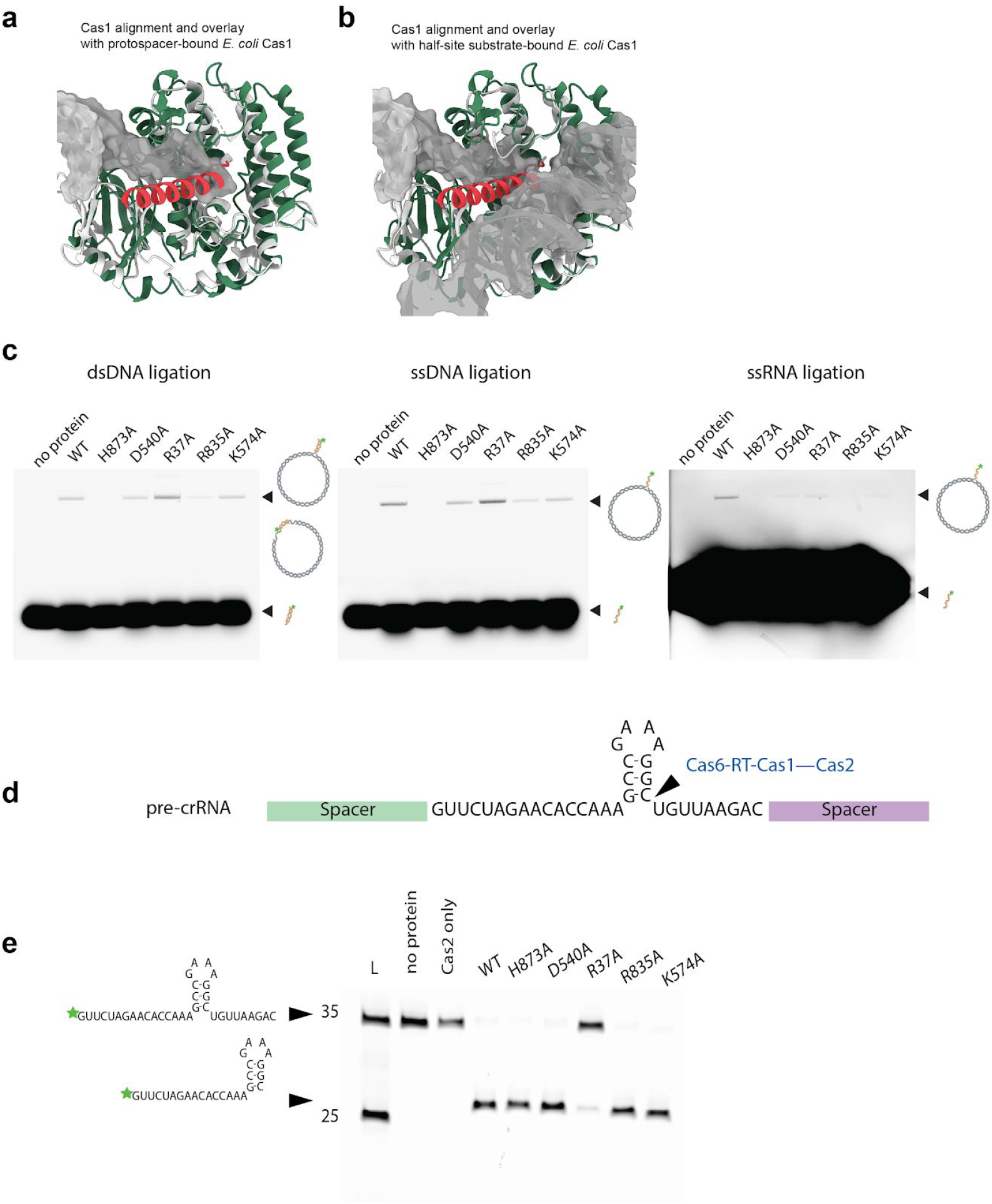
Crosstalk between Cas1, RT, and Cas6 domains. **a)** Alignment and overlay of Cas1a domain (dark green) with Cas1a from protospacer-bound *E. coli* Cas1—Cas2 structure (white, PDB: 5DS5). Protospacer substrate shown in gray surface representation. **b)** Alignment and overlay of Cas1a domain (dark green) with Cas1a from half-site substrate-bound *E. coli* Cas1—Cas2 structure (white, PDB: 5VVJ). Half-site substrate shown in gray surface representation. **c)** Ligation of fluorescent 35-nt dsDNA (0.5 M), ssDNA (0.5 M), and ssRNA (4 M) protospacer into target pCRISPR by WT and mutant Cas6-RT-Cas1—Cas2s. Integration products and free protospacer are indicated and schematized. Star indicates 6-carboxyfluorescein label. **d)** Sequence and predicted secondary structure of repeat with predicted cleavage site indicated. **e)** crRNA processing activity assay comparing WT and mutant Cas6-RT-Cas1—Cas2s. Sequence and predicted secondary structure of CRISPR repeat RNA and the predicted product are indicated. Star indicates 6-carboxyfluorescein label. Uncropped gels are available in Supplementary Data Set 2.

**Supplementary Video 1**. 3D Variability analysis of cryo-EM data heterogeneity in Cas6-RT-Cas1-Cas2 complex.

**Supplementary Table 1.**
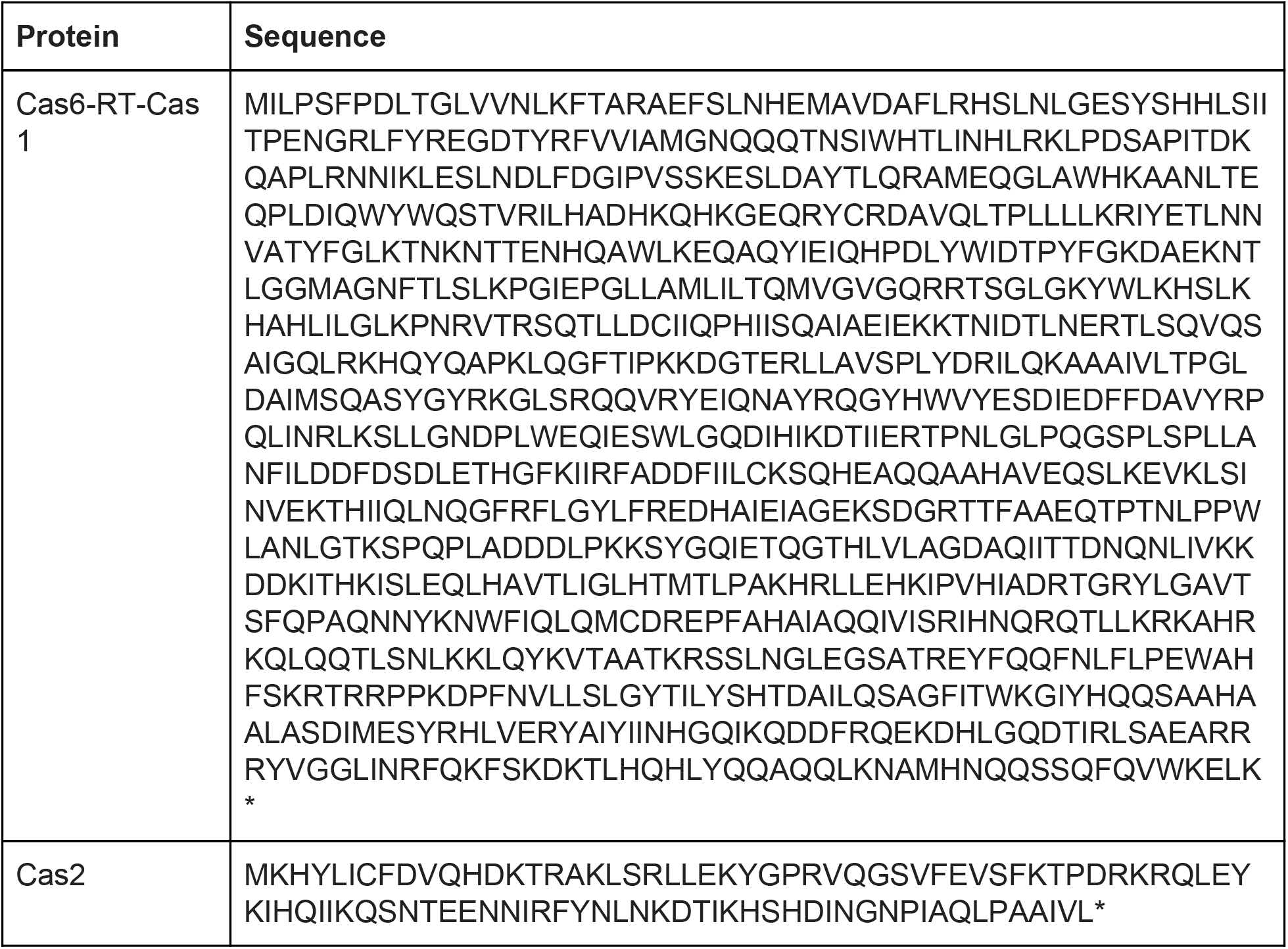
Sequences of Type III *Thiomicrospira* Cas6-RT-Cas1 and Cas2 Proteins.

**Supplementary Table 2.**
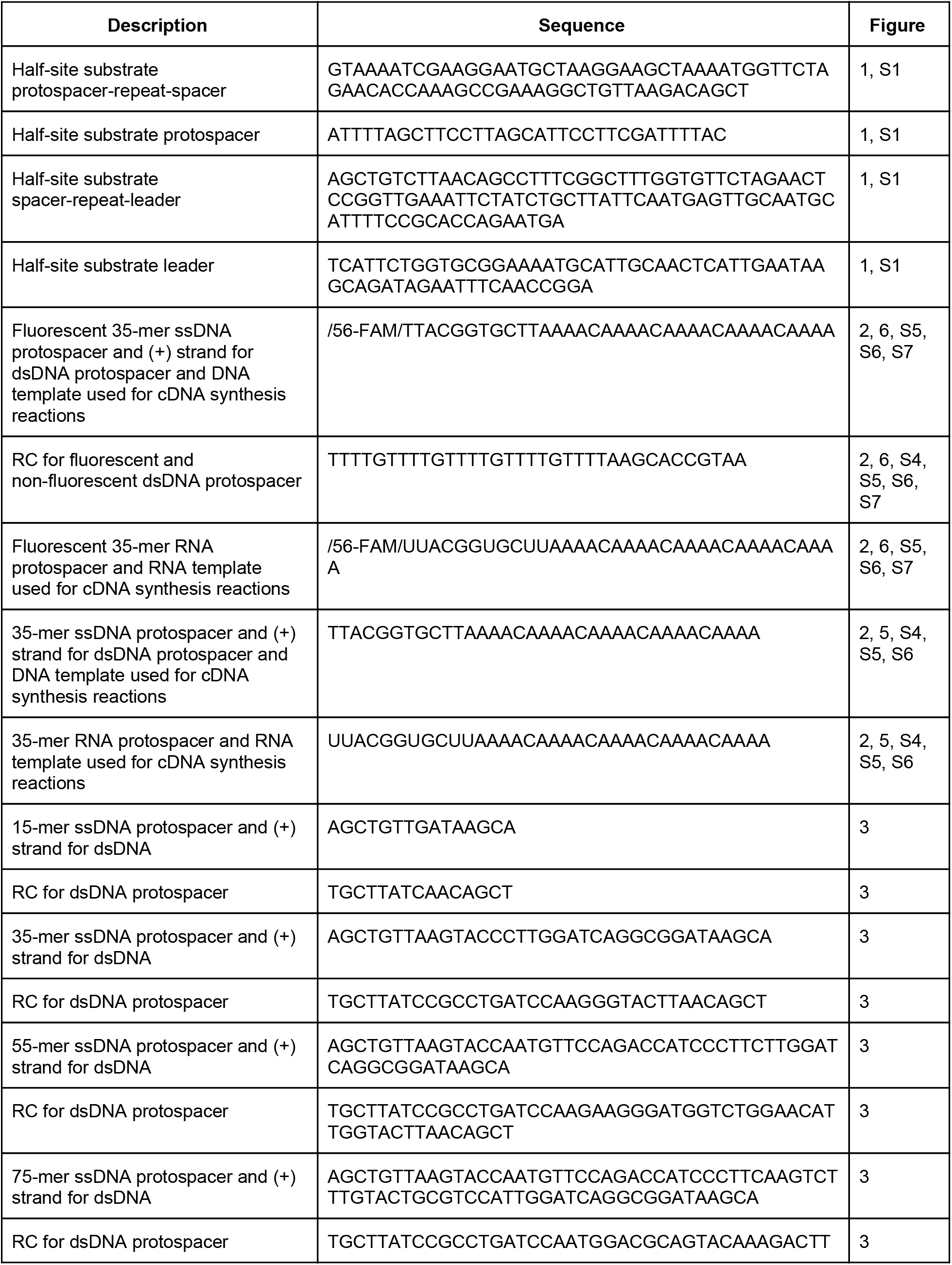
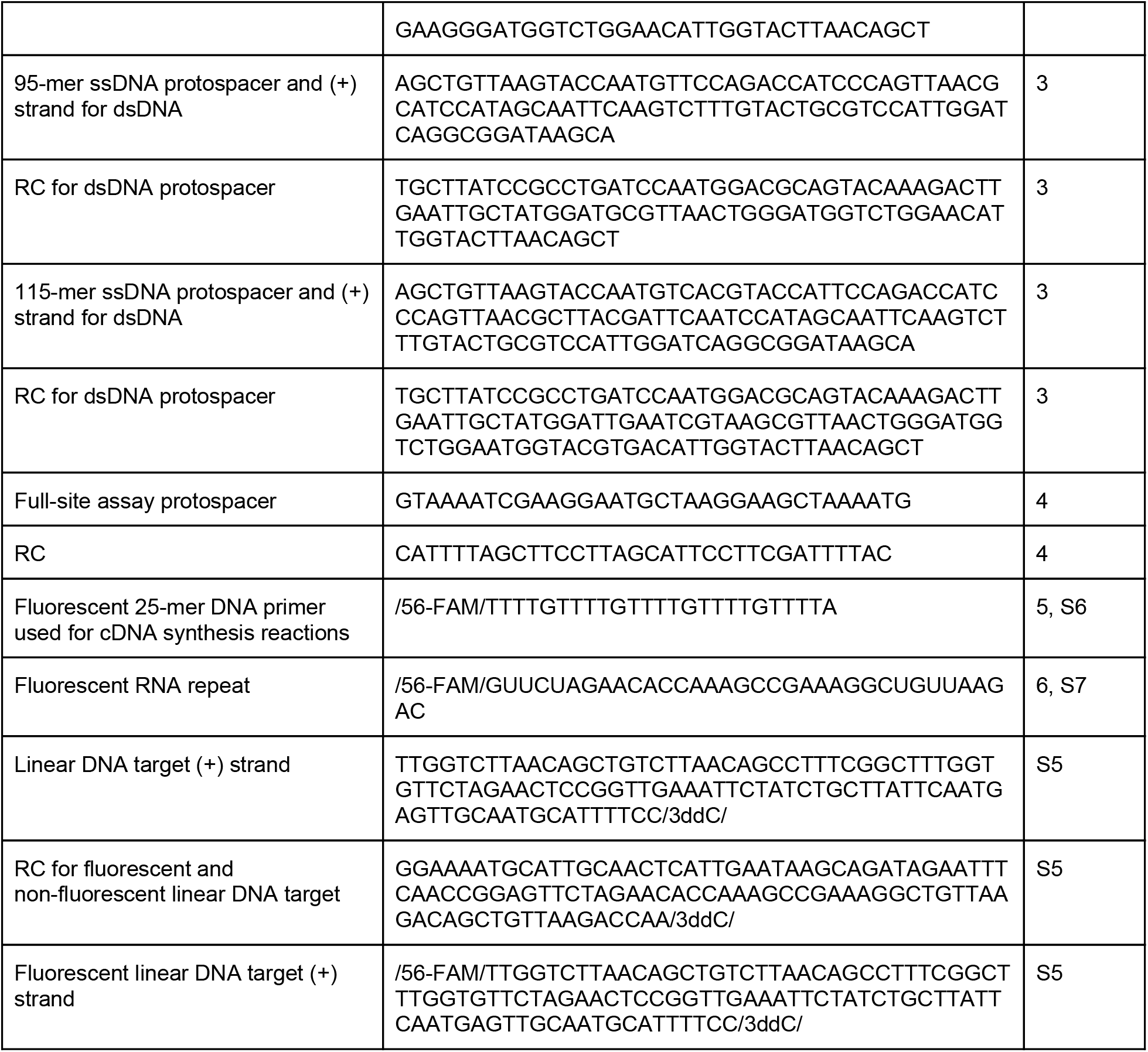
DNA substrates used in this study. RC indicates the complementary strand of the previous oligonucleotide. Sequences are written 5’ to 3’.

**Supplementary Table 3.**
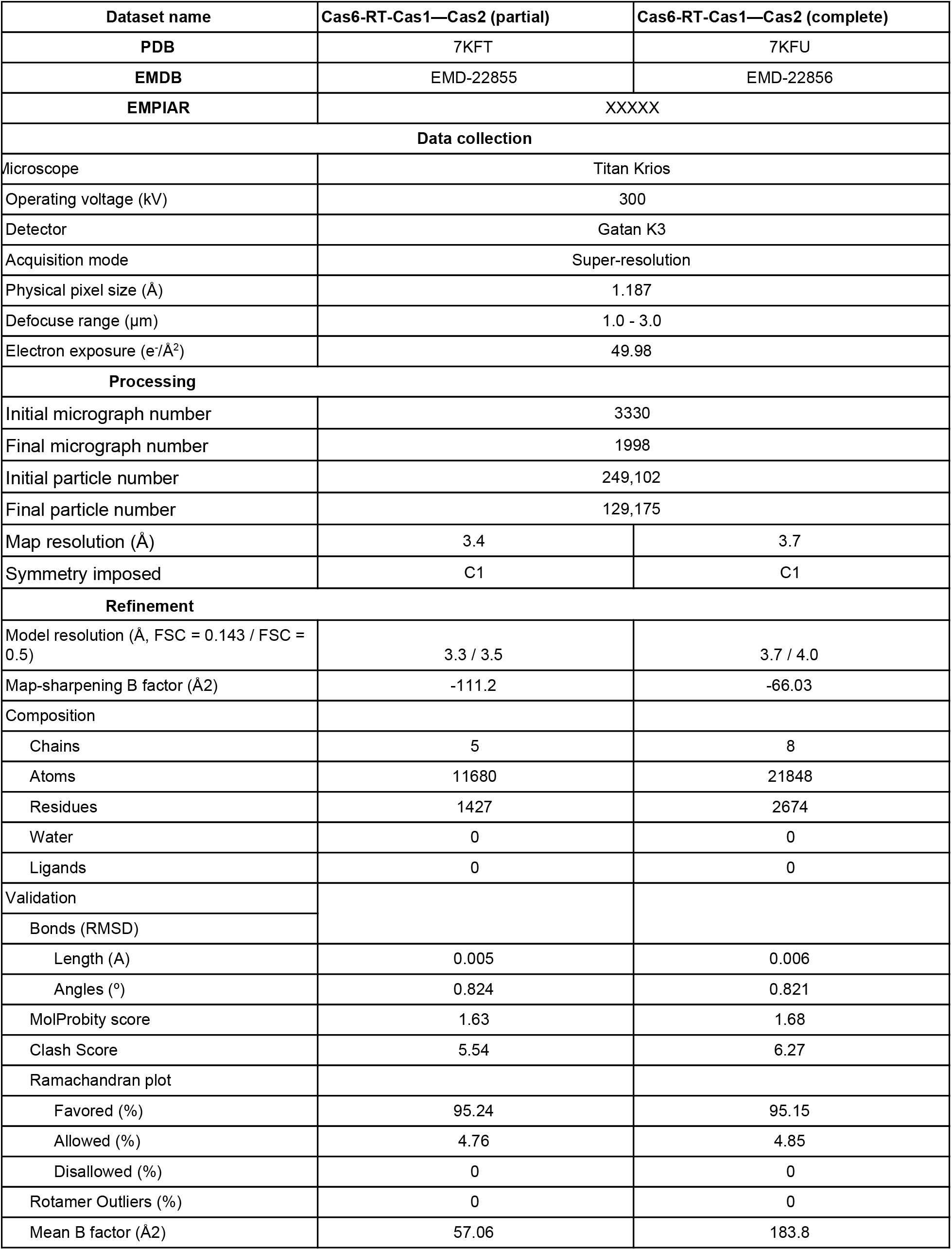
Cryo-EM data collection, processing, and refinement.

