## Supplementary Data Set 2 for "Structural coordination between active sites of a Cas6-reverse transcriptase-Cas1—Cas2 CRISPR integrase complex"

### Uncropped Gels

Figure 2b

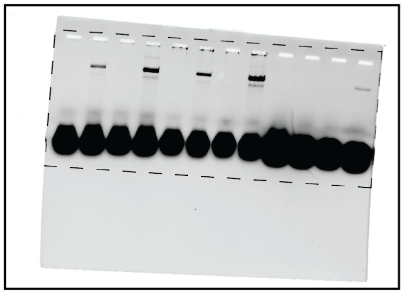

Figure 3c

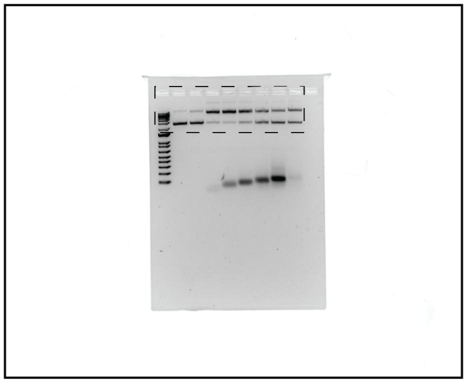

Figure 3d

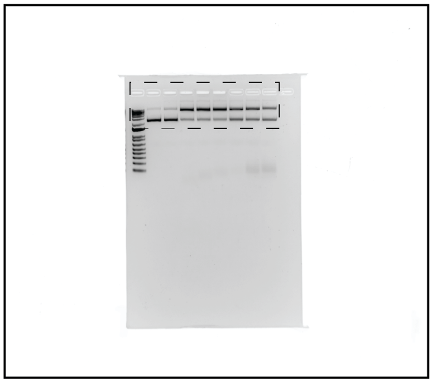

Figure 5e

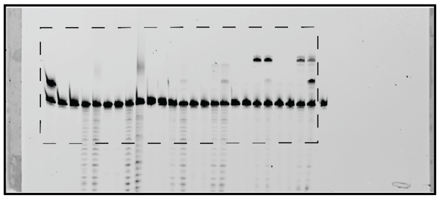

Figure 5f

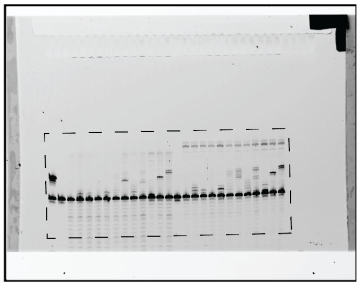

Supplementary Figure 4a dsDNA

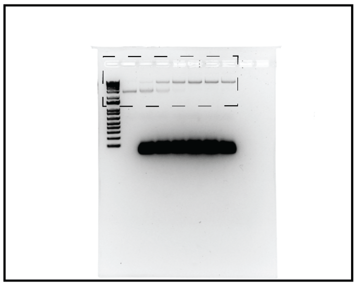

Supplementary Figure 4a ssDNA

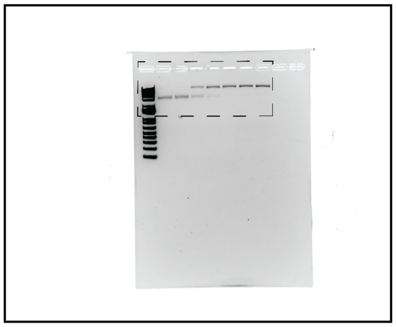

Supplementary Figure 4a ssRNA

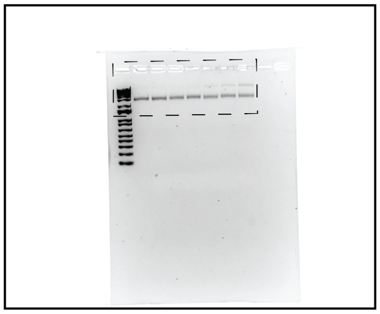

Supplementary Figure 4a no substrate

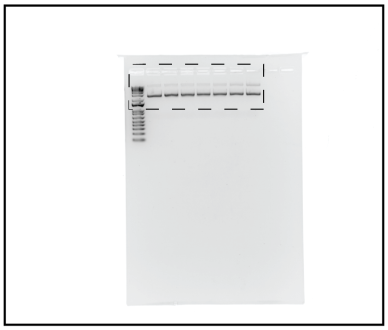

Supplementary Figure 5b.
Assay gel, fluorescent

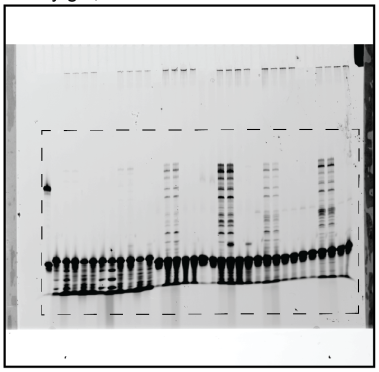

Supplementary Figure 5b.
Ladder, SYBR Gold post-stained

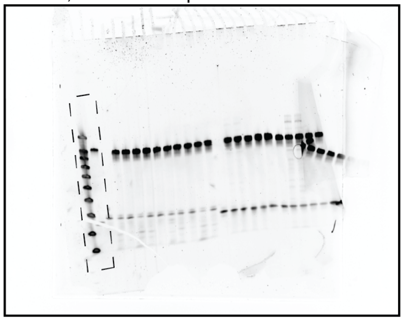

Supplementary Figure 5c.
Assay gel, fluorescent

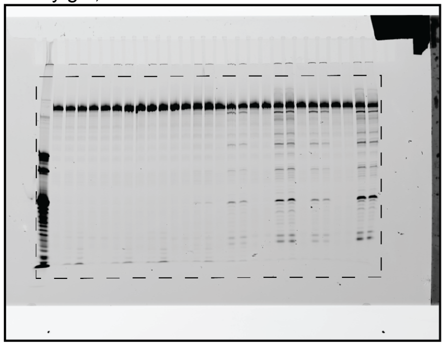

Supplementary Figure 5c.
Ladder, SYBR Gold post-stained

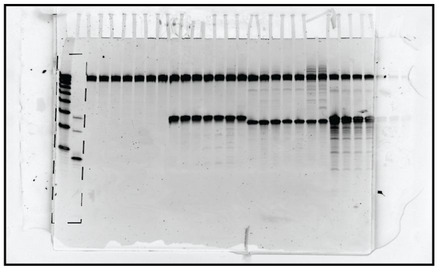

Supplementary Figure 6a.
Assay gel, fluorescent

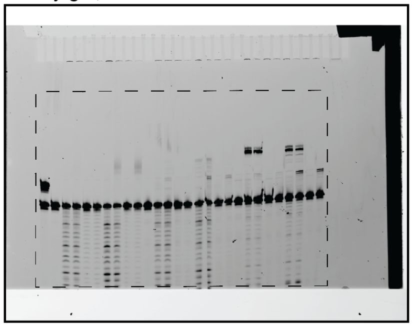

Supplementary Figure 6a.
Ladder, SYBR Gold post-stained

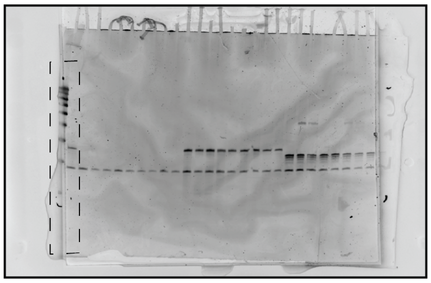

Supplementary Figure 6b.
Assay gel, fluorescent

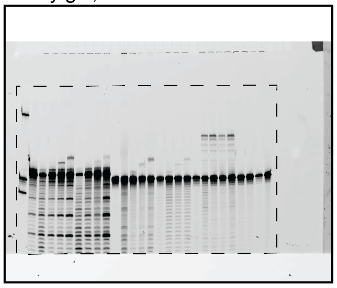

Supplementary Figure 6b.
Ladder, SYBR Gold post-stained

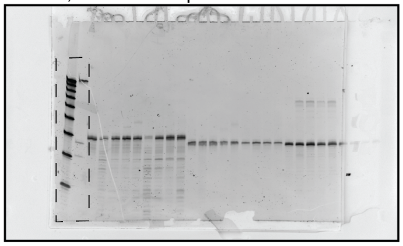

Supplementary Figure 7c. dsDNA ligation

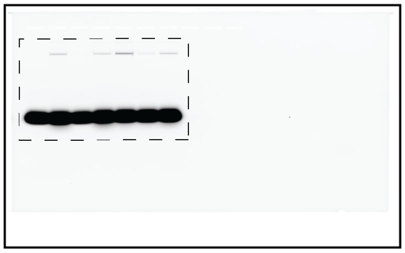

Supplementary Figure 7c.ssDNA ligation

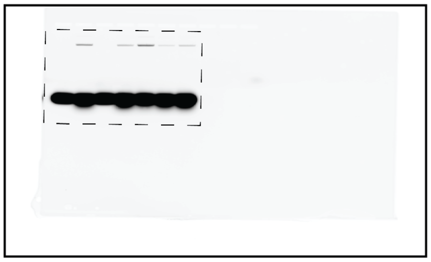

Supplementary Figure 7c.
ssRNA ligation

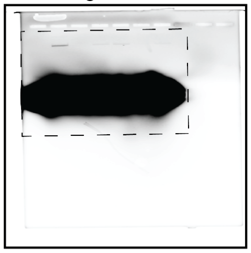

Supplementary Figure 7e

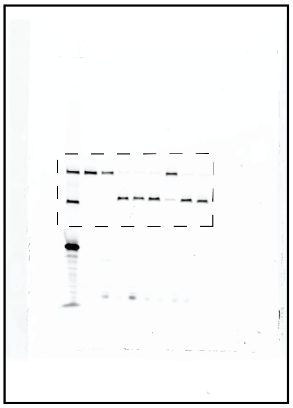
